# Single-cell intracellular pH dynamics regulate the cell cycle by timing G1 exit and the G2 transition

**DOI:** 10.1101/2021.06.04.447151

**Authors:** Julia S. Spear, Katharine A. White

**Affiliations:** Department of Chemistry and Biochemistry, University of Notre Dame, Notre Dame, IN, USA; Harper Cancer Research Institute, University of Notre Dame, Notre Dame, IN, USA

**Keywords:** Cell Cycle, Intracellular pH, Single-Cell Methods

## Abstract

Transient changes in intracellular pH (pHi) regulate normal cell behaviors, but roles for spatiotemporal pHi dynamics in single-cell behaviors remains unclear. Here, we mapped single-cell spatiotemporal pHi dynamics during mammalian cell cycle progression both with and without cell cycle synchronization. We found that single-cell pHi is dynamic throughout the cell cycle: pHi decreases at G1/S, increases in mid-S, decreases at late S, increases at G2/M, and rapidly decreases during mitosis. Importantly, while pHi is highly dynamic in dividing cells, non-dividing cells have attenuated pHi dynamics. Using two independent pHi manipulation methods, we found that low pHi inhibits completion of S phase while increased pHi promotes both S/G2 and G2/M transitions. Our data also suggest that low pHi cues G1 exit, with decreased pHi shortening G1 and increased pHi elongating G1. Furthermore, dynamic pHi is required for S phase timing, as high pHi elongates S phase and low pHi inhibits S/G2 transition. This work reveals spatiotemporal pHi dynamics are necessary for cell cycle progression at multiple phase transitions in single human cells.

## INTRODUCTION

In normal epithelial cells, intracellular pH (pHi) is near neutral (∼7.2) while extracellular pH (pHe) is more alkaline (∼7.4) (White et al., 2017a). Transient changes in pHi driven by ion transporter activity (Boron, 2004) have been shown to regulate normal cell behaviors such as differentiation (Ulmschneider et al., 2016), proliferation (Flinck et al., 2018b), migration (Choi et al., 2013; Martin et al., 2011), and apoptosis (Sergeeva et al., 2017). However, most studies of pHi-dependent cell behaviors are limited because average pHi is measured across a population of cells, pHi measurements are performed in non-native cellular environments, or pHi is monitored over short timeframes during a long biological process. Thus, our understanding of how spatiotemporal single-cell pHi dynamics regulate cell behaviors is limited. Better understanding of how pHi dynamics drive single-cell behaviors will reveal mechanistic roles for pHi in regulating biology and validate pHi as a reporter of cell phenotype.

One pHi-dependent behavior where rigorous spatiotemporal single-cell pHi measurements could help reveal mechanism is cellular proliferation. Links between cell cycle and pHi were first identified in unicellular organisms, such as tetrahymena (Gillies and Deamer, 1979), *Dictyostelium* (Aerts et al., 1985), and *S. pombe* (Karagiannis and Young, 2001). In tetrahymena, two increases in pHi (∼0.4 pH units) were observed pre- and post-S phase (Gillies and Deamer, 1979). However, all measurements of pHi changes were made at the population level and used harsh synchronization techniques (starvation and heat shock) that can disrupt essential cell metabolic functions in addition to regulating the cell cycle (Gillies and Deamer, 1979). In *Dictyostelium*, increased pHi (∼0.2 pH units) was measured during S phase and when pHi was artificially increased, DNA replication and protein synthesis were increased (Aerts et al., 1985). However, no timing or delays in S phase progression were noted with pHi manipulation, and only population-level pHi measurements were reported. In conflict with these two previous studies, no relationship between cell cycle progression and pHi was found in *S. pombe* when pHi was monitored using a genetically-encoded pHi biosensor (pHluorin) (Karagiannis and Young, 2001). Therefore, the existing data in unicellular organisms is inconsistent on whether pHi dynamics are sufficient to regulate (or time) cell cycle progression. We note that these inconsistencies could be biologically meaningful and due to species-specific differences in cell cycle regulation, or the inconsistencies could be artifactual (due to non-physiological pHi measurements and manipulations in these models).

Some studies in animal cell models have also shown a relationship between pH and cell cycle progression. In quiescent populations of human tumor cells, it was shown that a narrow range of pHe values (pH 6.8 to 7.2) are required to recruit cells into the cell cycle (Taylor and Hodson, 1984). While this suggests that a defined range of pHe is required for normal proliferation, the authors did not measure pHi during these experiments (Deutsch et al., 1982). Population-level analyses of pHi in thymidine-synchronized MCF-7 breast cancer cells showed that pHi fluctuated after thymidine release but no statistical significance was noted (Flinck et al., 2018a). Strengthening the link between pHi and cell cycle regulation, knockdown of the Na^+^-H^+^ exchanger (NHE1) and the Na^+^-HCO_3_^−^transporter (NBCn1) caused elongation of S phase and a delay in the G2/M transition in breast cancer cells (Flinck et al., 2018a), but single-cell pHi was not measured. In another example, an increase in pHi driven by NHE1 was found to be required for G2/M transition in fibroblasts, but single-cell pHi was not measured and pH was manipulated using genetic knockout or overexpression of NHE1 (Putney and Barber, 2003). As ion transporters also serve scaffolding and signaling roles, genetic knockdown produces transport-independent effects on cell biology. In summary, although these studies lay a strong framework for a relationship between pHi and cell cycle, single-cell pHi measurements in mammalian cells cultured under native environments are needed to elucidate how temporal pHi dynamics regulate cell cycle progression.

Here, we measure single-cell pHi under physiological growth conditions in both asynchronous and synchronized cell populations to determine how pHi regulates cell cycle progression in single cells. We found that single-cell pHi oscillates during cell cycle progression. Importantly, we determined that pHi oscillations correlate with cell cycle stages: pHi decreases near the G1/S transition, increases during mid-S, decreases again at S/G2 transitions, and finally increases at G2/M followed by rapid acidification during mitosis. Using pHi manipulation and Fluorescent Ubiquitination-based Cell Cycle Indicator (FUCCI) reporters, we determined that dynamic pHi is necessary for normal cell cycle progression. Similar to prior work, we show that increased pHi is required for successful completion of G2/M. But our work also reveals previously uncharacterized pHi dynamics regulating both G1 exit and S phase duration. This work highlights advantages of using single-cell pHi measurements to investigate single-cell behaviors like cell cycle progression and suggests mechanisms for limiting pHi-dependent cell cycle progression in diseases with dysregulated pH, such as cancer (increased pHi) (Harguindey et al., 2017; White et al., 2017a) and neurodegeneration (decreased pHi) (Majdi et al., 2016).

## RESULTS

### Single-cell pHi in clonal cell lines is heterogeneous

We first examined whether asynchronous single-cell pHi measurements under physiological conditions could recapitulate population-level averages while also reporting on physiological single-cell heterogeneity. We used a genetically-encoded pH biosensor, mCherry-pHluorin (mCh-pHl) (Koivusalo et al., 2010), which has been used to measure pHi in cultured cells (Choi et al., 2013; Koivusalo et al., 2010) and tissues (Grillo-Hill et al., 2015) and has a dynamic linear range between pH 6.5 and 8.0 (Grillo-Hill et al., 2014). Briefly, direct measurement of pHi in single living cells can be achieved by performing ratiometric imaging of pHluorin/mCherry fluorescence intensities. Fluorescence of pHluorin is pH-sensitive in the physiological range while mCherry fluorescence is pH-insensitive and used to normalize for biosensor expression. At the end of the experiment, single-cell standardization is performed using isotonic buffers of known pH containing the protonophore nigericin (Fig. 1A). This method of pHi measurement avoids issues of uneven dye loading, washout, and photobleaching associated with pH-sensitive dyes (Grillo-Hill et al., 2014).

**Figure 1:**
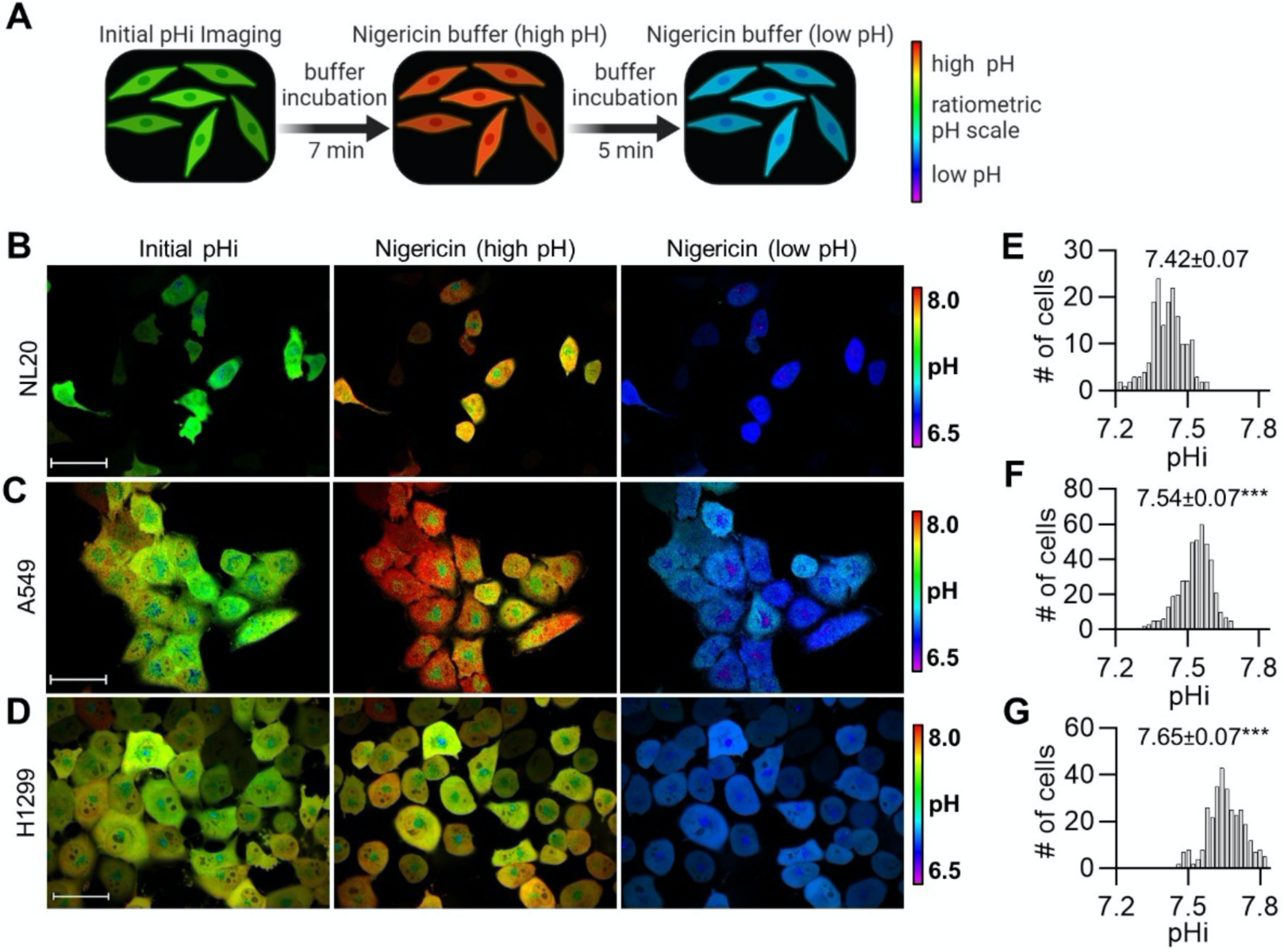
Intracellular pH is heterogeneous in normal and cancerous lung cell lines and median pHi significantly increases in cancer cells. A) Schematic of single-cell pHi measurements using a stably expressed pH biosensor, mCherry-pHluorin (mCh-pHl), and the protonophore nigericin to standardize the biosensor (see methods for details). B-D) Representative images of pHi measurements and standardization in B) NL20, C) A549, and D) H1299 cells stably expressing mCh-pHl. Ratiometric display of pHluorin/mCherry fluorescence ratios, scale bars: 50 μm. E-G) Histograms of single-cell pHi in E) NL20 (n=173, 3 biological replicates), F) A549 (n=424, 4 biological replicates), and G) H1299 (n=315, 3 biological replicates). Histograms are binned at 0.02 pH units. Above histograms, median±S.D. is shown. Significance was determined by a Mann Whitney test. (***p<0.001, compared to NL20).

We stably expressed mCh-pHl in normal lung epithelial cells (NL20), primary tumor site-derived lung cancer cells (A549), and metastatic site-derived lung cancer cells (H1299). We chose these lung-derived cells because these clonal cell lines are well characterized in literature, are morphologically heterogeneous, and tolerate stable expression of the mCh-pHl biosensor. We first confirmed that biosensor expression does not alter pHi homeostasis in these cells by comparing population pHi measurements of the clonal biosensor lines (NL20-mCh-pHl, A549-mCh-pHl and H1299-mCh-pHl) to matched parental cell lines (Grillo-Hill et al., 2014) (Fig. S1A-C). One distinct advantage of single-cell imaging experiments for pHi measurement is that pHi can be measured directly in conditioned media without the need to use fresh bicarbonate-or HEPES-based isotonic washing solutions that are required for population level assays (see methods for solution composition). Thus, single-cell pHi measurements are more likely to reflect accurate pHi setpoints and dynamics of cells and give better comparison to other cell biological assays or signaling profiles measured from cells cultured continuously in complete media.

We next measured single-cell pHi in individual NL20-mCh-pHl (Fig. 1B), A549-mCh-pHl (Fig. 1C), and H1299-mCh-pHl (Fig. 1D) cells. Representative pHluorin and mCherry channels and single-cell standardization curves can be found in Fig. S1D-I. To assay pHi heterogeneity in these clonal cell lines, we prepared distribution histograms of single-cell pHi measurements and found that the pHi of primary tumor cells (A549-mCh-pHl) (Fig. 1F; 7.54±0.07) was increased compared to normal lung epithelial cells (NL20-mCh-pHl) (Fig. 1E; 7.42±0.07). Importantly, metastatic tumor cells (H1299-mCh-pHl) had the highest median pHi (Fig. 1G; 7.65±0.07), which was significantly higher than both the normal and primary tumor clonal cell lines. To support these results, we also measured single-cell pHi in metastatic triple negative breast cancer cells (MDA-MB-231) and found it was also significantly increased (7.52±0.18) compared to matched normal breast epithelial cells (MCF10A) (7.23±0.11) (Figure S2). Taken together, we find that aggressive cancer cell lines have higher single-cell pHi compared to normal epithelial cells across multiple tissue origins. Importantly, our data show that pooled single-cell pHi measurements reveal significant heterogeneous pHi distributions that are lost in population-level analyses. These data also suggest that even genetically-identical clonal cell lines exhibit single-cell pHi heterogeneity that may be biologically meaningful and could report on non-genetic cell phenotype such as cell cycle status.

### Cells released from G1 synchronization have dynamic pHi

Next, we sought to measure pHi dynamics during cell cycle progression. We synchronized H1299-mCh-pHl cells using Palbociclib, which blocks phosphorylation of the retinoblastoma protein and synchronizes cells prior to the G1 checkpoint (Liu et al., 2018) (Fig. 2A). Palbociclib is an efficient G1 synchronizer in H1299 cells, with nearly 85% synchronization after 24 h treatment and minimal DNA damage (Trotter and Hagan, 2020). After Palbociclib synchronization, cells were imaged at 0, 4, 8, 12, 24, and 36 hours (h) after release (Fig. 2B) and single-cell pHi distributions were measured (Fig. 2C). Qualitatively, we noticed that cells were larger at earlier time points (0-4 h) and that at 12 h cells had cell rounding and smaller apparent cell size indicative of mitotic cells (Fig. 2B). We observed oscillating pHi distributions during cell cycle progression where single-cell pHi significantly decreased between 0 and 4 h, significantly increased between 4 and 8 h, decreased again between 8 and 12 h, and finally increased between 12 and 24 h (Fig. 2D). These data suggest that pHi is dynamic during cell cycle progression with temporally regulated fluctuations in pHi after synchronization release.

**Figure 2:**
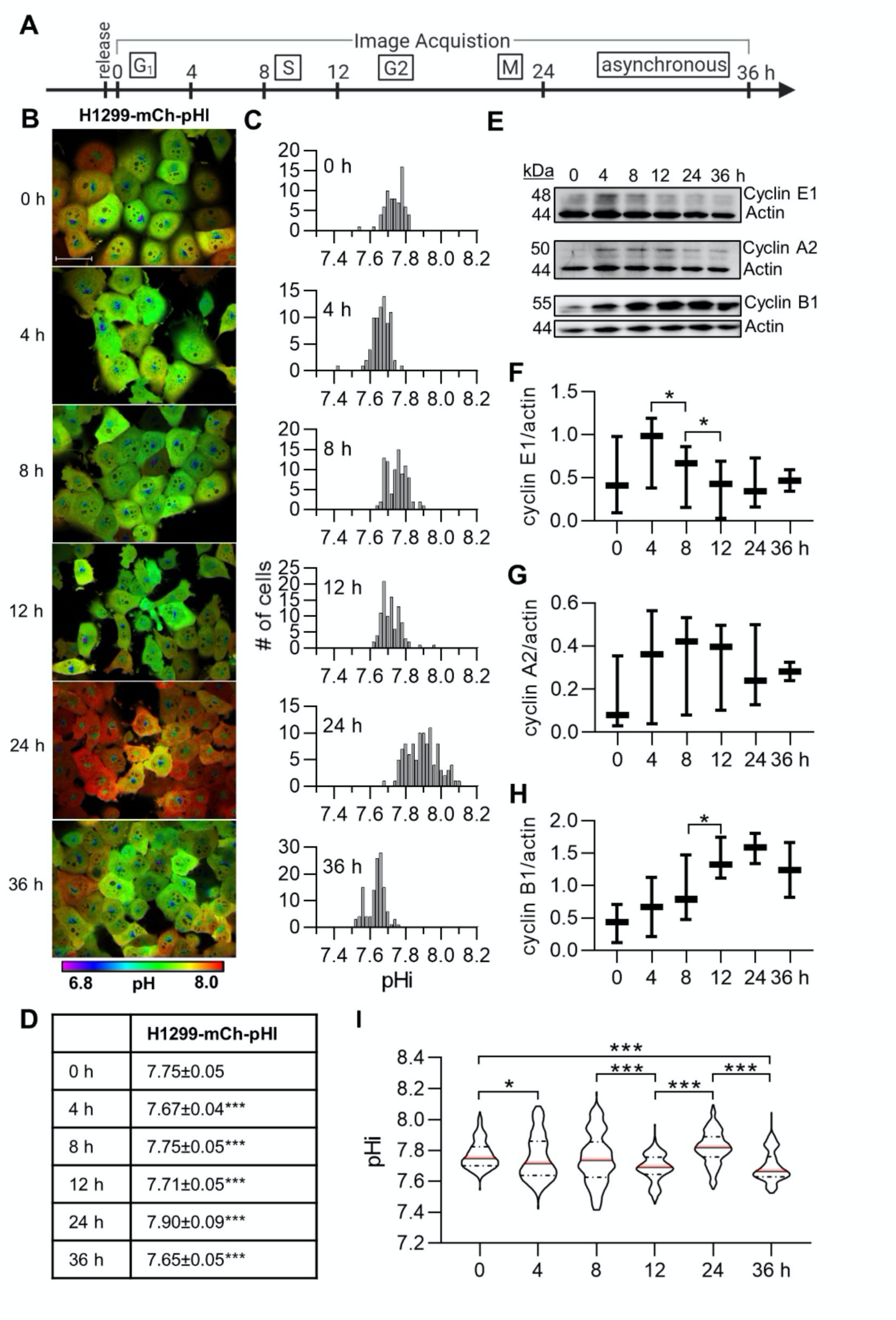
Intracellular pH is dynamic following G1 synchronization and correlates with cyclin levels. A) Schematic of image acquisition after Palbociclib synchronization. B) Representative images of H1299-mCh-pHl cells at indicated time points after release. Ratiometric display of pHluorin/mCherry fluorescence ratios. Scale bar: 50 μm. C) Histograms of single-cell pHi data collected as in B, from one biological replicate. Histograms binned at 0.02 pH units. Additional replicates in Fig. S3. D) Table of pHi values from data in C (median±S.D.). E) Representative immunoblots for cyclin E1, A2, and B1 with actin loading controls. Box and whisker plots of F) cyclin E1, G) cyclin A2, and H) cyclin B1 immunoblot data (3 biological replicates). Additional replicates in Fig. S3. I) Violin plots of raw pHi (median red bars; 0 h, n=231; 4 h, n= 253; 8 h, n=262; 12 h, n=273; 24 h, n=338; 36 h, n=262; 3 biological replicates). In D and I, significance was determined by Kruskal Wallis test with Dunn’s multiple comparisons correction. In F-H, significance was determined by paired t-test. In D and F-I, each time point was compared to the preceding time point and in I, 0 h was compared to 24 h. (*p<0.05; ***p<0.001).

We confirmed that Palbociclib appropriately synchronized the cells by immunoblotting for cyclins from matched cell lysates (Fig. 2E). Cyclin E1 regulates G1/S (Siu et al., 2012), cyclin A2 regulates S and G2 phases (De Boer et al., 2008), while cyclin B1 regulates G2 and must be degraded prior to anaphase in mitosis (Chang et al., 2003). We observe that cyclin E1, a marker of G1/S, significantly increases from 0 to 4 h, which is expected for a cell population properly synchronized in G1 phase (Fig. 2F). These cells undergo mitosis approximately 24 h post-Palbociclib release because cyclin B1 levels, an inducer of G2/M, peaked 24 h post-release (Fig. 2H). By 36 h, protein abundance was similar across all cyclins, as expected in an asynchronous population. Cyclin immunoblots and pHi agreed across 3 biological replicates (additional pHi replicates and blots in Fig. S3).

Single-cell pHi measurements on Palbociclib-treated cells were compared over three biological replicates and we found that pHi significantly decreased at the G1/S transition (4 h, 7.75±0.15) and in late S phase (12 h, 7.69±0.09), significantly increased at G2/M (24 h, 7.82±0.11), and then significantly decreased again at the end of the experiment in asynchronous cells (36 h, 7.67±0.10) (Fig. 2I). To assess if Palbociclib treatment alters resting pHi, we pooled synchronized time points (Fig. 2I, 0-24 h) and compared these data to pHi measurements at experiment endpoint (Fig. 2I, 36 h) and to untreated, asynchronous H1299-mCh-pHl pHi data (Fig. 1G). We note that pHi in Palbociclib-treated cells was significantly increased compared to untreated, asynchronous cells, indicating that Palbociclib synchronization may also alter pHi homeostasis (Fig. S3C). Previous work did find that Palbociclib induced markers of senescence and autophagy when used for >36 h (Capparelli et al., 2012), so this is a confounding factor on pHi at 24 h of treatment. However, the increases in resting pHi were uniform in our data and trends in pHi dynamics were robust across multiple biological replicates.

### Cells exhibit cell cycle-linked pHi dynamics independent of cell cycle synchronization method

To confirm that the temporal pHi dynamics observed in Figure 2 were linked to specific cell cycle phases and not an artifact, or off-target effect, of Palbociclib synchronization, we next synchronized H1299-mCh-pHl cells in early S phase using a double-thymidine block (Chen and Deng, 2018). Thymidine acts as a DNA synthesis inhibitor by accumulating dTTP and depleting dCTP within the cell (Bjursell and Reichard, 1973; Bolderson et al., 2004). We synchronized H1299-mCh-pHl cells and imaged them at 0, 4, 8, 12, and 24 h after thymidine release (Fig. 3A-B). Qualitatively, cells were larger at 0 h and by 8 h had altered morphology that could indicate mitosis (Fig. 3B). In this representative replicate, single-cell pHi significantly decreased between 0 and 4 h, significantly increased between 4 and 8 h, decreased between 8 and 12 h, and increased again between 12 and 24 h (Fig. 3C-D). This general trend supports pHi data from cells released earlier in the cell cycle (Palbociclib, G1) (Fig. 2). Importantly, the observed phase-shifted pHi oscillations confirm that pHi dynamics are linked to cell cycle timing and not experiment timing, regardless of the synchronization method used.

**Figure 3:**
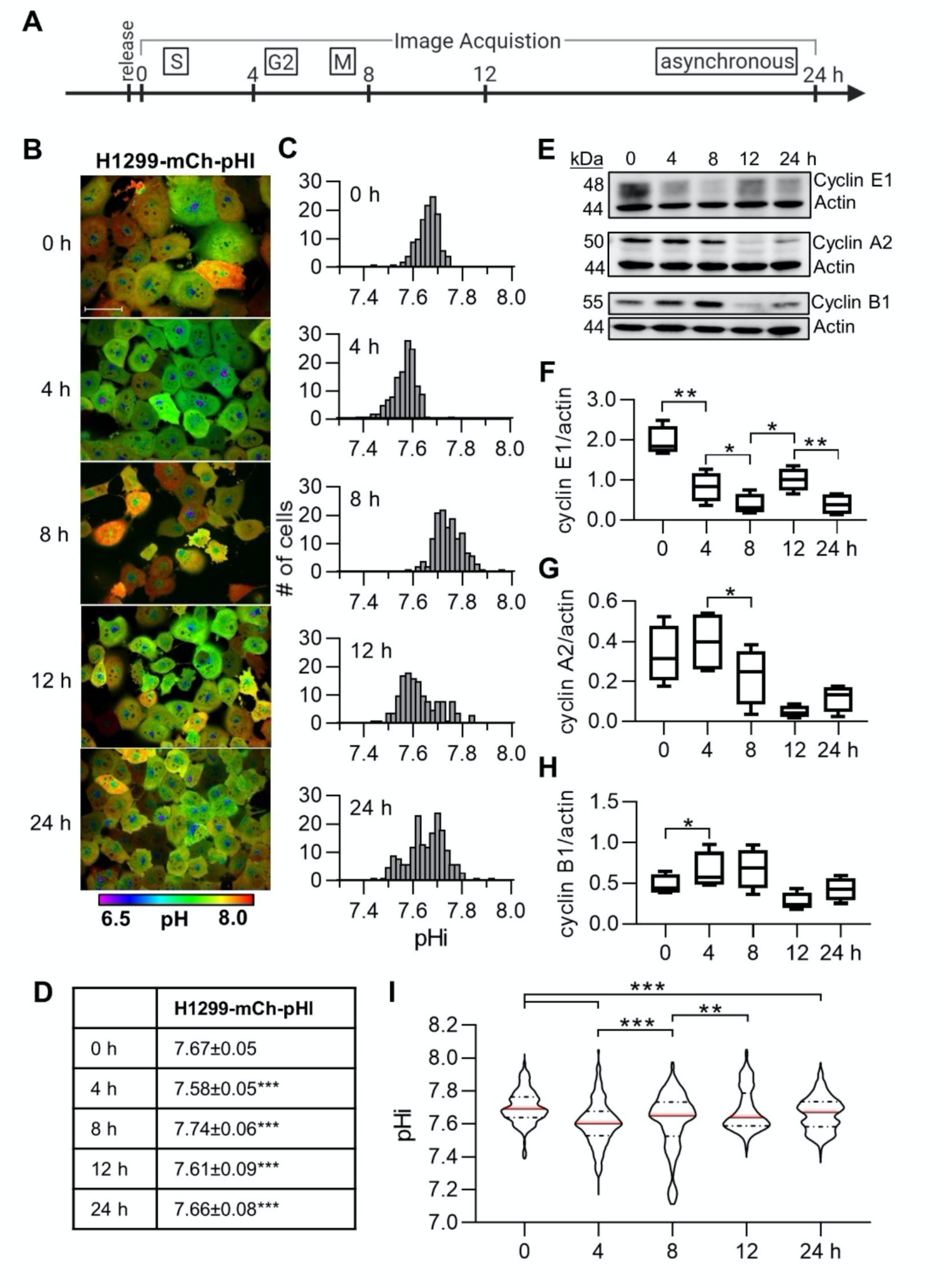
Intracellular pH is dynamic after release from early S phase in H1299-mCh-pHl cells and correlates with cyclin levels. A) Schematic of image acquisition after a double-thymidine synchronization. B) Representative images of H1299-mCh-pHl cells at indicated time points after release. Ratiometric display of pHluorin/mCherry fluorescence ratios, scale bar: 50 μm. C) Histograms of single-cell pHi data collected in B, from one biological replicate. Histograms binned at 0.02 pH units. Additional replicates in Fig. S4. D) Table of pHi values from data in C (median±S.D.). E) Representative immunoblots for cyclin E1, A2, and B1 with respective actin loading controls. Box and whisker plots of F) cyclin E1, G) cyclin A2, and H) cyclin B1 immunoblot data (4 biological replicates). Additional replicates in Fig. S5. I) Violin plots of raw pHi values (median red bars; 0 h, n=500; 4 h, n= 468; 8 h, n=517; 12 h, n=558; 24 h, n=652; 4 biological replicates). In D and I, significance was determined by a Kruskal Wallis test with Dunn’s multiple comparisons correction. In F-H, significance was determined by a paired t-test. In D and F-I, each time point was compared to its preceding time point and in I, 0 h compared to 24 h. (*p<0.05; **p<0.01; ***p<0.001).

We also confirmed that thymidine treatment appropriately synchronized the cells by immunoblotting for cyclins from matched cell lysates (Fig. 3E). We found that cyclin E1 (G1/S) peaks at 0 h, as expected for a cell population synchronized in early S phase (Fig. 3F). The cells undergo mitosis approximately 8 h after thymidine release as cyclin A2 (S/G2) was highest at 4 h and significantly decreased by 8 h (Fig. 3G), while cyclin B1 (G2/M) peaked 8 h after release (Fig. 3H). Cyclin E1 levels increased again by 12 h, suggesting that by 12 h most cells in this assay had completed the cell cycle and progressed back to G1 (Fig. 3F). By 24 h, protein abundance was similar across all cyclins, as we would expect in an asynchronous population. Immunoblots for additional replicates are shown in Fig. S4, and pooled cyclin results match previously published data on synchronized H1299 cell populations (Chen and Deng, 2018).

Single-cell pHi measurements from pooled thymidine-treated biological replicates revealed that the median pHi of the cell populations decreased significantly at 4 h (late S phase), increased from 4 to 8 h (G2/M), and decreased again from 8 to 12 h (M/G1) (Fig. 3I). This preserves the oscillating pHi pattern measured in the individual replicate (Fig. 3B-C, additional replicates in Fig. S4). Thymidine synchronization did not alter homeostatic pHi when compared to pooled synchronized thymidine data (Fig. 3I, 0-12 h), asynchronous thymidine data (Fig. 3I, 24 h), or untreated H1299-mCh-pHl cells (Fig. 3G) (Fig. S4C). The single-cell pHi data after thymidine release recapitulates both the decreased pHi in late S phase and increased pHi at G2/M that we measured after Palbociclib release, while also revealing a significant pHi increase during early S phase.

To confirm that cell-cycle linked pHi dynamics are not unique to H1299 cells, we synchronized A549-mCh-pHl with a double-thymidine block and observed similar pHi dynamics (Fig. S5). Cell morphology and pHi oscillations matched H1299-mCh-pHl at respective time points (Fig. S5A), with a decrease in pHi from 0 h to 4 h, an increase from 4 h to 8 h, and decreases at 12 h and 24 h (Fig. S5B-C). Again, synchronization was confirmed using cyclin immunoblots, where cyclin A2 (S/G2) peaked at 4 h, cyclin B1 (G2/M) peaked at 8 h, and both proteins were low at 12 h indicating the start of a new cycle with cells in G1 (Fig. S5D). Like H1299-mCh-pHl cells, pooled A549-mCh-pHl cell pHi significantly increased from 4 h to 8 h (G2/M) and decreased following mitosis at 12 h (Fig. S5E). Observing identical cell-cycle linked pHi dynamics across different cell models suggests pHi increases leading to division (G2/M, 4-8 h) and decreases after division (G1, 8-12 h) may be necessary for division timing and re-entry into the cell cycle.

From these data, we conclude that pHi is dynamic through the cell cycle at the single-cell level: pHi decreases during G1/S, increases in early S phase, decreases leading to S/G2, increases prior to G2/M, and decreases following mitosis.

### Single cells alkalize prior to G2/M, followed by rapid acidification during mitosis and pHi recovery in daughter cells

In the previous experiments, cells from matched populations were identically treated and released from synchronization for imaging at various time points after release. These time points showed that single-cell pHi distributions oscillate with cell cycle progression, but the snapshots may not reflect continuous single-cell pHi dynamics or cell cycle progression phenotypes. To address this limitation, we established a time-lapse microscopy approach to track pHi dynamics over an entire cell cycle in a single cell.

We first measured pHi changes in single asynchronous H1299-mCh-pHl cells. We observed cells randomly dividing throughout the time-lapse, indicating the cells were asynchronous (Fig. S6A), and representative stills of ratiometric time-lapse pHi imaging in a dividing cell are shown (Fig. 4A, Movie 1). For this cell, pHi quantification shows oscillating pHi dynamics similar to those observed in the snapshot experiments (Figure 4B). Although we were able to mark entry into mitosis via DNA condensation at prophase, we cannot determine other cell phase transitions with this approach. We noted there is a prominent alkalization in the hours prior to mitosis (Figure 4A-B, 19 h), followed by rapid acidification during mitosis (Figure 4A-B, P-C). To compare trends in single-cell pHi dynamics across many individual cells, we selected prophase as a “normalization point” for each individual dividing cell. We observed a significant period of alkalization that began ∼7 h prior to division and persisted until prophase (Fig. 4C). These pHi increases in single dividing cells correlate with the increased pHi observed during G2/M in the discontinuous endpoint data (Fig. 2I, 3I, S5E) and suggest that increased pHi may be a required signal for division of single cells. The single-cell time-lapse analysis also allows us to distinguish cells that undergo mitosis and cells that do not. If pHi dynamics are a sufficient regulator of cell cycle progression, we may expect to see attenuated pHi dynamics in non-dividing cells. To test this hypothesis, we quantified pHi from non-dividing cells. Representative stills of ratiometric time-lapse pHi imaging in a non-dividing cell from the asynchronous population are shown (Fig. 4D, Movie 2). We found that pHi dynamics are attenuated in non-dividing cells compared to dividing cells (Fig. 4E-F), suggesting that pHi dynamics are correlated with successful cell cycle progression. Thus, pHi dynamics may be an important biomarker for or driver of normal cell cycle progression.

**Figure 4:**
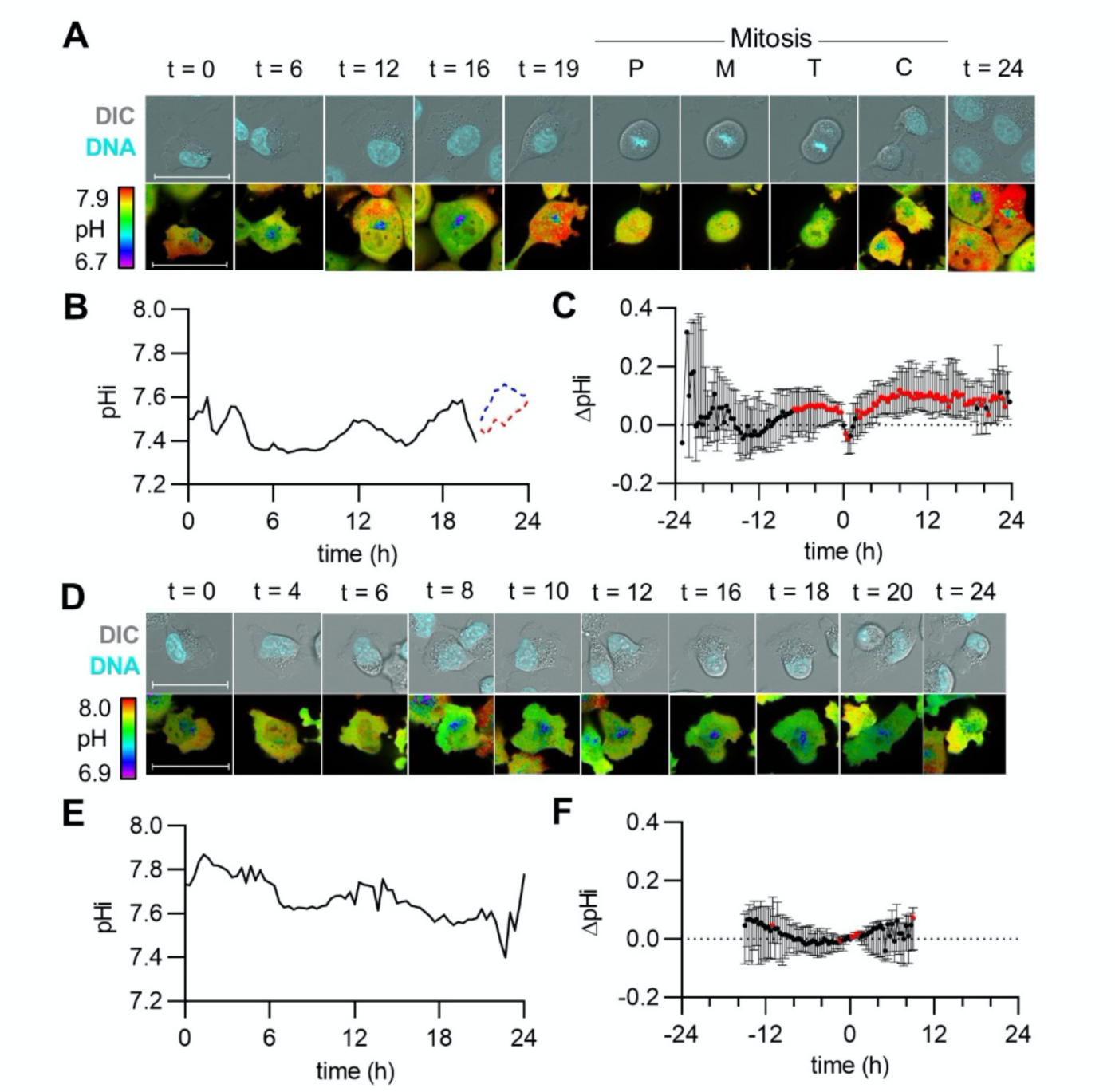
Intracellular pH increases leading to G2/M, followed by rapid acidification prior to division and pHi recovery in daughter cells. A) Representative stills from Movie 1 of a dividing H1299-mCh-pHl cell at indicated time (h). Top is Hoechst dye (DNA, cyan) and DIC merge. Bottom is ratiometric display of pHluorin/mCherry fluorescence ratios, scale bars: 50 μm. B) Traces of calculated pHi values of the cell in A) (black, solid line) and in daughter cells (red and blue dotted lines). C) pHi changes in dividing cells, relative to pHi at prophase (P) for each individual cell (median±IQ range, n=39, 4 biological replicates). Significance was determined by a one-sample Wilcoxon test compared to 0 (red points, *p<0.05). D) Representative stills from Movie 2 of a non-dividing H1299-mCh-pHl cell at indicated time (h). Top is Hoechst dye (DNA, cyan) and DIC merge. Bottom is ratiometric display of pHluorin/mCherry fluorescence ratios, scale bars: 50 μm. F) Trace of pHi values of cell in E) (black, solid line) over time. G) pHi changes in non-dividing cells, relative pHi at 15 h (median±IQ range, n=25, 4 biological replicates). Significance was determined by a one-sample Wilcoxon test compared to 0 (red points, *p<0.05).

In order to directly compare single-cell time-lapse pHi dynamics to the prior data, we next collected time-lapse pHi measurements in thymidine-synchronized H1299-mCh-pHl cells. We observed bursts of mitotic cells at 15 h, indicating thymidine was appropriately synchronizing individual cells (Fig. S6B), and representative stills of ratiometric time-lapse pHi imaging in a dividing cell are shown (Fig. 5A, Movie 3). For this dividing cell, pHi increased through late S and G2 phases (matched to Fig. 3 cyclin timing data), decreased during mitosis, and recovered rapidly in daughter cells (Fig. 5B). Similarly to the asynchronous time-lapse data above, we used prophase as a “normalization point” for comparing pHi in individual dividing cells and observed oscillating pHi dynamics with a significant period of alkalization beginning ∼11 h prior to division and persisting until prophase, followed by a rapid acidification during mitosis and recovery in daughter cells (Fig. 5C). Non-dividing cells had significantly attenuated pHi dynamics compared to dividers at the single-cell level (Fig. 5D-E, Movie 4). To compare trends in single-cell pHi dynamics for non-dividing cells, change in pHi was calculated for each non-dividing cell from the average prophase time in the synchronized time-lapses (15 h). We again observed attenuated pHi dynamics in single non-dividing cells (Fig. 5F). Importantly, the extended alkalization observed prior to prophase (Fig. 4C, Fig. 5C) is not observed in either non-divider dataset. This suggests that dynamic pHi could be a hallmark for cells moving through the cell cycle.

**Figure 5:**
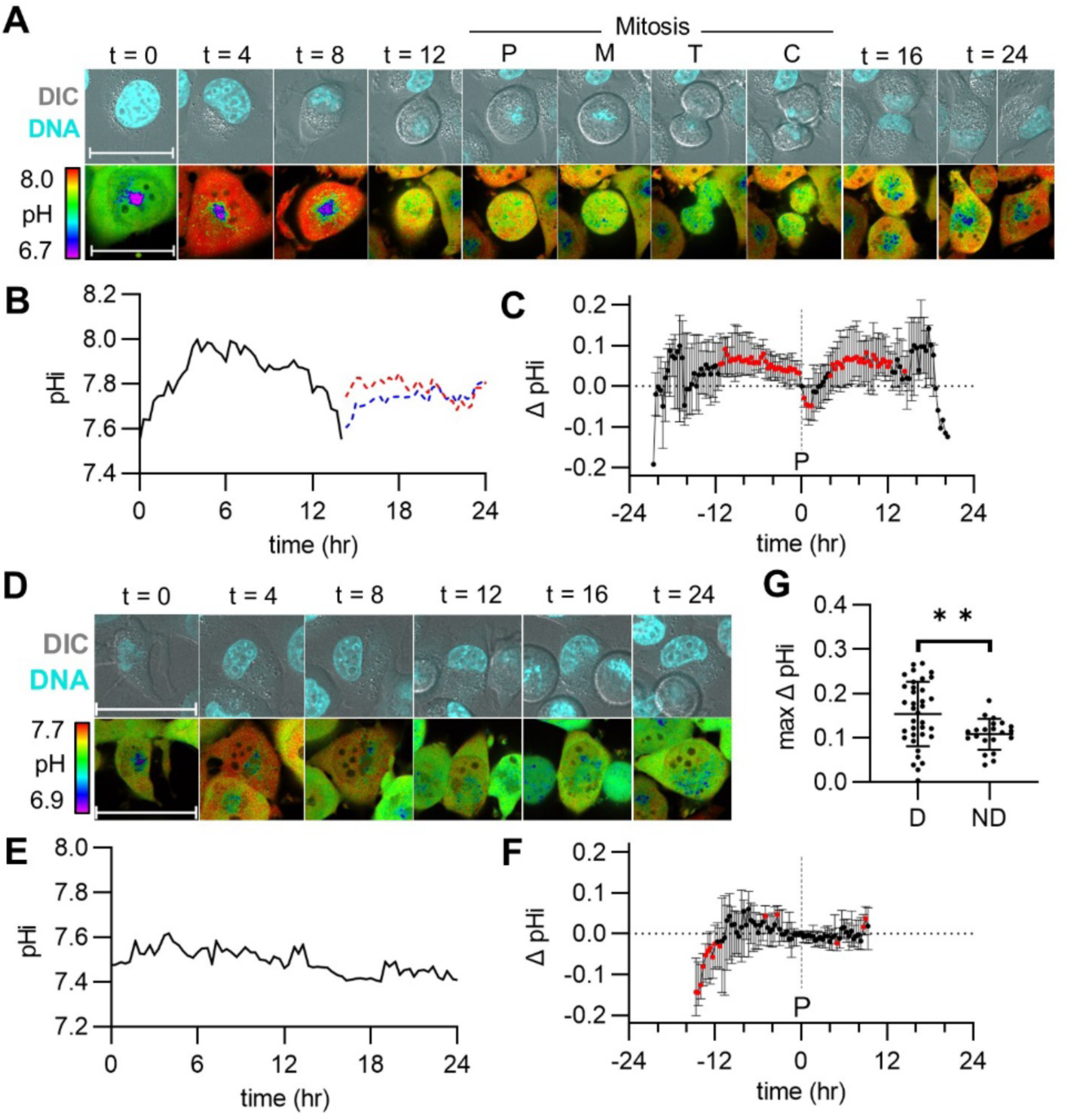
Cells released from S phase synchronization show pHi increases leading to G2/M, rapid acidification prior to division, and pHi recovery of daughter cells. A) Representative stills from Movie 3 of a dividing H1299-mCh-pHl cell at indicated time (h). Top is Hoechst dye (DNA, cyan) and DIC merge. Bottom is ratiometric display of pHluorin/mCherry fluorescence ratios, scale bars: 50 μm. B) Traces of calculated pHi values of the cell in A) (black, solid line) and in daughter cells (red and blue dotted lines). C) pHi changes in dividing cells, relative to pHi at prophase (P) for each individual cell (median±IQ range, n=39, 3 biological replicates). Significance was determined by a one-sample Wilcoxon test compared to 0 (red points, *p<0.05). D) Representative stills from Movie 4 of a non-dividing H1299-mCh-pHl cell at indicated time (h). Top is Hoechst dye (DNA, cyan) and DIC merge. Bottom is ratiometric display of pHluorin/mCherry fluorescence ratios, scale bars: 50 μm. E) Trace of pHi values of cell in D (black, solid line) over time. F) pHi changes in non-dividing cells, relative to the pHi at average time of prophase (determined from dividing cell data) (median±IQ range, n=22, 3 biological replicates). G) Scatter plot of max pHi change in individual dividing (D) and non-dividing (ND) cells (mean±S.D.). For D and F, significance was determined by a one-sample Wilcoxon test compared to 0 (red points, *p<0.05). For G, significance determined by an unpaired t-test (**p<0.01).

Since cell synchronization confers increased consistency of cell phase transitions, we binned the thymidine data depending on timing of prophase (Fig. S6B). Importantly, we observed significant alkalizations in each group ∼5 hours prior to prophase regardless of mitosis timing (Fig. S6C), which correlates with G2 entry based on previous data in H1299 cells (Rajal et al., 2021) and our own data with FUCCI cell cycle reporters (shown below; G2 is ∼4.0 h). These data strongly suggest that the increased pHi we observe across all time-lapse datasets at 5 h coincides with G2 entry. Next, to confirm that the dynamics we observed in the time-lapse are not an artifact of increased mCh-pHl biosensor expression or altered biosensor photobleaching rates, we tracked mCherry and pHluorin intensities over time in both dividers and non-dividers from the synchronous time-lapses (Fig. S6D-E). We observed that mCherry fluorescence dynamics in dividing cells show similar trends to non-dividing cells across the time-lapse experiment (Fig. S6D). Furthermore, the pHluorin increases observed over time in dividing cells are not correlated with increased mCherry fluorescence, indicating observed pHluorin increases are not due to increases in biosensor expression (Fig. S6E) but instead reflect dynamic pHi in single cells.

These time-lapse data suggest that increased single-cell pHi dynamics may be correlated with (or regulate) single-cell cell cycle progression. Supporting this hypothesis, parent cells that divided within the 24 h period showed a significantly higher median pHi increase (0.16±0.07; Fig. 5G) when compared to non-dividing cells (0.10±0.03; Fig. 5G). We note that the magnitude of pHi changes observed in single dividing cells (∼0.16 pH units) corresponds well with physiological pHi increases previously reported in single cells during other cell behaviors, such as cell migration (0.1-0.35) (Denker and Barber, 2002).

Taken together, these time-lapse data suggest that single dividing cells have oscillating pHi dynamics with an increase in pHi leading to mitosis, a rapid acidification during mitosis, and recovered pHi in daughter cells. Our next question was whether pHi dynamics regulate or time cell cycle progression.

### Dysregulated pHi dynamics affect cell cycle phase duration and cause phase-specific arrests

We have shown pHi is dynamic and correlates with cell cycle phases in asynchronous cells, cells synchronized at G1 with Palbociclib (Fig. 2), and cells synchronized at early S phase with thymidine (Fig. 3). To compare pHi data from both synchronization techniques, we aligned pHi data (Fig. 2I, Fig 3I) according to significantly increased cyclin B1 expression (Fig. 2H, Fig. 3H) and found oscillating pHi dynamics throughout the cell cycle (Fig. 5A). We observed that pHi decreases during G1/S, increases in mid-S phase, decreases prior to S/G2, and increases prior to G2/M (Fig. 6A). In addition, the time-lapse data in asynchronous (Fig. 4C) and thymidine-synchronized single cells (Fig. 5C) confirmed the dynamic increases in pHi leading to G2/M and uncovered a rapid acidification during M phase (Fig. 6A). These data suggest a correlation between pHi and cell cycle progression, but to determine a causal relationship, we sought to manipulate pHi and monitor effects on cell cycle phases in real-time. To do this, we established pHi manipulation techniques (Larsen et al., 2012; White et al., 2017b) and used the FUCCI cell cycle reporter (Grant et al., 2018) to track single cells during cell cycle progression.

**Figure 6:**
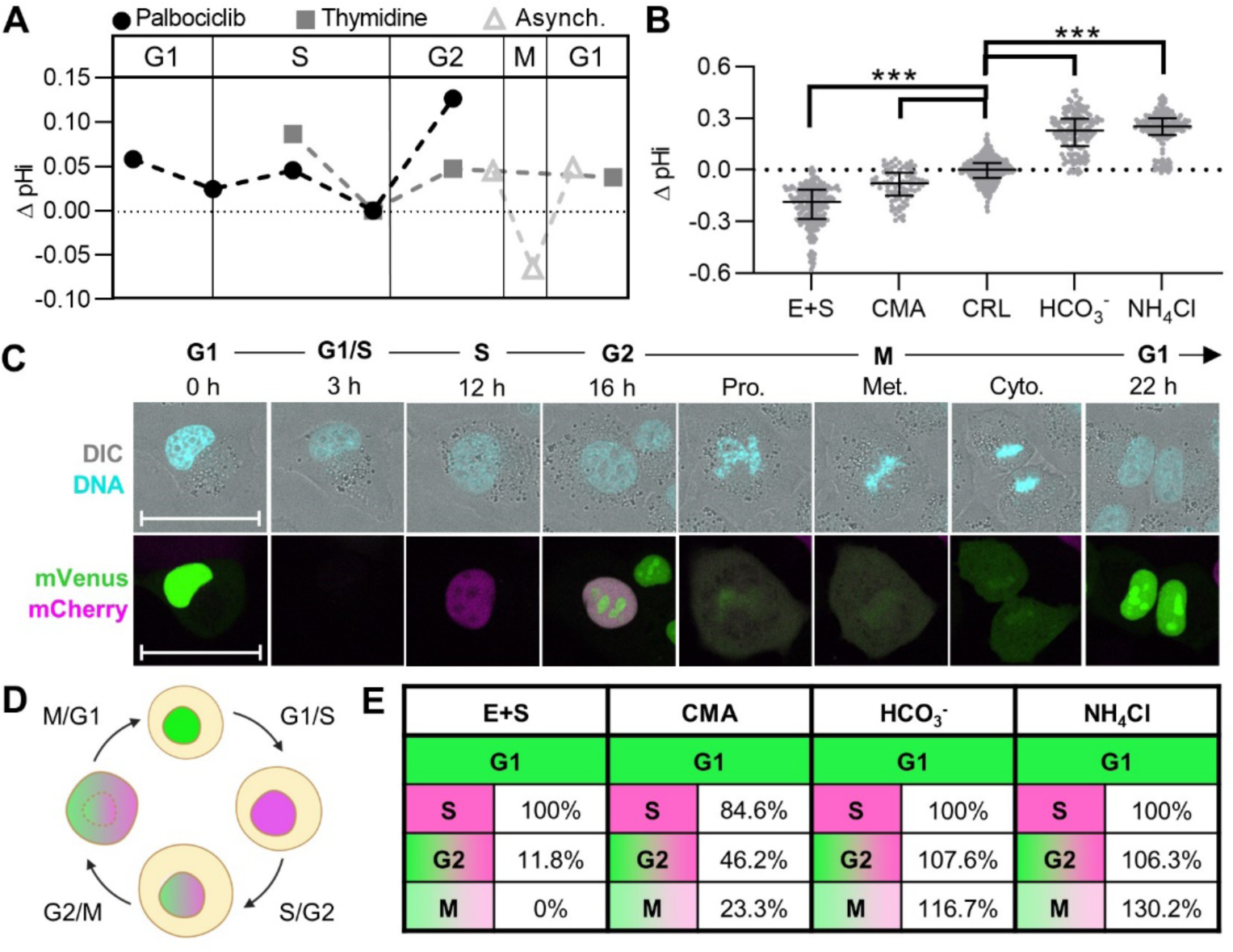
Single-cell pHi manipulation shows that pHi dynamics are key regulators of cell cycle. A) Median plots of single-cell delta pHi from synchronizations and asynchronous (Asynch.) time-lapses. Data reproduced from Fig. 2I, 3I, and 4C (thymidine, 4 h, n=4; Palbociclib, 12 h, n=3; Asynch., n=4) B) Single-cell pHi of H1299-mCh-pHl cells treated for 24 h with 15 μM EIPA and 30 μM S0859 (E+S, n=233) or 1 μM Concanamycin A (CMA, n=79) to lower pHi, untreated (CRL, n=602), or supplemented with 100 mM NaHCO_3_ (HCO_3_^−^, n=146) or 20 mM ammonium chloride (NH_4_Cl, n=193) to raise pHi (see methods for details). Additional treatment time points are shown in Fig. S8. C) Representative stills from Movie 5. Shown is a single H1299-FUCCI cell with PIP-mVenus (green) and mCherry-Geminin (magenta) tracked through each cell cycle phase. Hoechst dye (DNA, cyan) and DIC merge shown; scale bars: 50 μm. D) Schematic of PIP-FUCCI reporter fluorescence during cell cycle phase transitions (Grant et al., 2018). E) Successful phase entry of cells starting in G1, where each treatment is normalized to matched controls. E+S (n=27), CMA (n=13), HCO_3_^−^ (n=13), or NH_4_Cl (n=13). For B, scatter plots (median±IQ range), with Mann-Whitney test to determine statistical significance (***p<0.001).

To manipulate pHi, we used combinations of selective ion transporter inhibitors to lower pHi and media supplementation to raise pHi. To lower pHi, we used: Concanamycin A (CMA), which inhibits V-ATPases (Huss et al., 2002); 5-(N-ethyl-N-isopropyl)amiloride (EIPA), which inhibits NHE1 (White et al., 2017b); and 2-chloro-N-[[2′-[(cyanoamino)sulfonyl][1,1′-biphenyl]-4-yl]methyl]-N-[(4-methylphenyl)methyl]-benzamide (S0859) (Larsen et al., 2012), which inhibits the Na^+^-HCO_3_^−^ transporter (NBCn1) (see methods for details). Both incubation with CMA and combination treatment with EIPA and S0859 (E+S) lowered pHi compared to CRL (lowering pHi by ∼0.08 and ∼0.18 pH units, respectively; Fig. 6B). To raise pHi, we supplemented the media with ammonium chloride (NH_4_Cl) or with bicarbonate (HCO_3_^−^) (raising pHi by ∼0.23 and ∼0.25 pH units compared to CRL, respectively; Fig. 6B).

FUCCI reporters use regulatory domains of cell cycle proteins to differentially express fluorescent proteins and report on cell cycle progression in single cells (Fig. 6C-D). We used the PIP-FUCCI reporter, which allows improved delineation of S phase (Grant et al., 2018) compared to older FUCCI variants. PIP-FUCCI reporter fluorescence is driven by regulatory domains of PCNA interacting protein degron from human Cdt1_1-17_ (PIP) fused to mVenus and Geminin_1-110_ fused to mCherry. PIP-mVenus accumulates in the nucleus during G1 and is rapidly lost during onset of DNA replication (S phase). At the beginning of S phase, mCherry-Geminin accumulates and is expressed throughout S, G2, and M phases. During the S-G2 transition, mVenus accumulates again, and both mVenus and mCherry are co-expressed until division. Thus, the PIP-FUCCI reporter system enables accurate delineation of both G1/S and S/G2. M phase is marked by nuclear envelope breakdown and diffusion of mVenus and mCherry fluorescent proteins throughout the cell. Mitosis and cytokinesis can also be monitored by DNA stain and DIC imaging (Fig. 6C). Following cytokinesis, only mVenus is expressed in the two daughter cell nuclei, marking G1.

To determine how pHi dynamics regulate cell cycle progression, we stably expressed PIP-FUCCI in H1299 cells (H1299-FUCCI) (Fig. 6C, Movie 5) and applied the validated pHi manipulation techniques to experimentally raise and lower pHi in cells (Fig. 6B). Using time-lapse confocal microscopy, we tracked single cells (with and without pHi manipulation) over a 36 h period and analyzed mVenus and mCherry fluorescent intensities to determine successful progression of single-cell cell cycle phases (see methods for details).

First, we analyzed the successful transition of cells from G1 to subsequent cell cycle phases in each pHi manipulation treatment compared to untreated (CRL) cells. We observed some common effects of pHi manipulation on the ability of single cells to progress normally through the cell cycle. First, successful S/G2 transitions were reduced when pHi was lowered with either E+S (11.8%) or CMA (46.2%) and increased when pHi was raised with either HCO_3_^−^ (107.6%) or NH_4_Cl (106.3%) (all compared to CRL, Fig. 6E). Second, successful G2/M transitions were similarly reduced when pHi was lowered with E+S (0%) or CMA (23.3%) and increased when pHi was raised with either HCO_3_^−^ (116.7%) or NH_4_Cl (130.2%) (all compared to CRL, Fig. 6E). We also note that the phenotype of successful phase transitions correlated with the magnitude of pHi changes with low pHi manipulation: larger decreases in pHi induced by E+S produced stronger phenotypes compared to smaller pHi decreases induced by CMA. These data indicate that pH dynamics do regulate successful cell cycle phase transitions, where decreased pHi is detrimental to S/G2 and G2/M transitions and increased pHi promotes these transitions in single cells.

We next wanted to explore how pHi manipulation alters the length of cell cycle phases. We first tested whether the selected pHi manipulation approaches induce replicative stress, as it has been previously shown that replication stress can cause cell cycle phase dysregulation (Matthews et al.; Técher et al., 2017). We treated cells with each of the pHi manipulation techniques (Fig. 6B) and stained the cells for gamma-H2AX (H2AX), a common marker of replicative stress (Kuo and Yang, 2008). We found that only CMA significantly increased H2AX staining, so we removed that treatment condition from further analysis (Fig. S7). While prior work has shown that high concentrations of EIPA can induce H2AX (Rolver et al., 2020), we did not observe induction of H2AX with E+S in our system. We next considered whether altered metabolism contributed significantly to the observed results. While supplementation with HCO_3_^−^ raised pHi (Fig. 6B), it also can alter extracellular pH (which we noted by change in phenol red indicator in the HCO_3_^−^-treated media) and also drastically alters cellular metabolism (Krycer et al., 2017; Li et al., 2020) which can lead to dysregulated cell cycle phases independent of pHi (Toshihiro Mitaka, 1991). For this reason, we also did not continue with full phase length quantification with HCO_3_^−^ treatment.

One benefit of using FUCCI reporters is that they can be used to assay cell cycle phase completion but can also be used to directly measure cell cycle phase length (Fig. 7A). To quantify cell cycle phase length, we plotted single-cell traces of FUCCI fluorescence intensities (mVenus, mCherry) aligned to division time (Fig. 7B-D). Importantly, we validated that even within 4 hours of treatment the pHi manipulations were sufficient to significantly change pHi (Fig. S8). Using fluorescence intensity cutoffs to determine G1/S and S/G2 (Fig. 7A, see methods), we measured significant differences in phase durations with high and low pHi conditions (Fig. 7E-H). From the single cell traces, we first noted that G1 phase in daughter cells (dotted lines, after 0 h) was altered in both E+S and NH_4_Cl treated cells compared to CRL. G1 phase (high mVenus, low mCherry) was significantly shortened in daughter cells at low pHi (Fig. 7E, 1.7±0.4 h) and significantly elongated at high pHi (Fig. 7E, I 8.7±2.3 h) compared to CRL cells (Fig. 7E, 6.0±1.8 h). These pHi-dependent changes in G1 phase duration indicate that low pHi may be a cue for G1 exit and that aberrant alkalization delays this cell cycle transition. These data align with the measured decreased pHi at G1/S in the prior endpoint assays (Fig. 6A).

**Figure 7:**
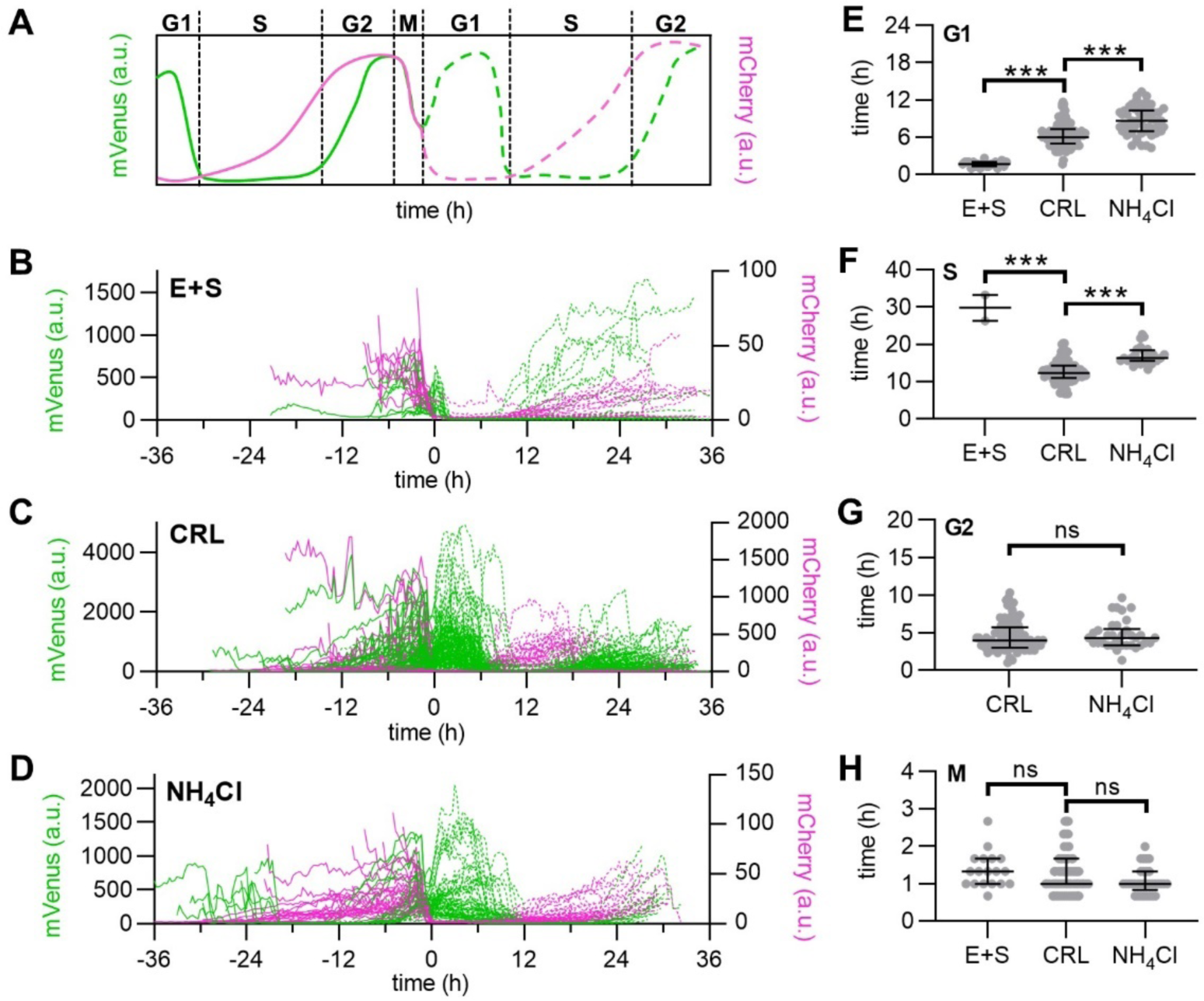
Single-cell FUCCI traces show low pHi is a cue for G1 exit, S phase requires high and low pHi, and S/G2 requires high pHi. A) Schematic of PIP-mVenus (green) and mCherry-Geminin (magenta) fluorescence intensities during cell cycle phases. B-D) Traces from single H1299-FUCCI cells treated as in Fig. 6B (E+S: 15 μM EIPA + 30 μM S0859; NH_4_Cl: 20 mM NH_4_Cl). Traces aligned at time of division at 0 h and daughter cells are indicated by dotted lines: E+S, (n=23); C) CRL (n=187); D) NH_4_Cl (n=72). (CRL and NH_4_Cl: 3 biological replicates, E+S, 2 biological replicates). E-H) Cell cycle phase durations from all cell populations (dividers and non-dividers). E) G1 (E+S, n=22; CRL, n=151; NH_4_Cl, n=51) F) S (E+S, n=3; CRL, n=88; NH_4_Cl, n=26), G) G2 (CRL, n=90; NH_4_Cl, n=34), and H) M (E+S, n=18; CRL, n=113; NH_4_Cl, n=33). For E-H, scatter plots (median±IQ range), with Mann-Whitney test to determine statistical significance (*p<0.05; ***p<0.001).

We also found that high pHi significantly elongated S phase (Fig. 7F, 16.3±2.4 h) compared to CRL (Fig. 7F, 12.3±3.0 h), while low pHi inhibited the S/G2 transition for all but 7.8% of cells (Fig. 7E). This suggests that high pHi is a requirement for S phase transition to G2, but there is also a need for low pHi for correct timing of S phase duration. The requirement for an increase and decrease in pHi is supported by the synchronized single-cell pHi data, which showed an increase in pHi during mid-S phase and a decrease in late S phase or near the S/G2 transition (Fig. 6A). Thus, our data suggests that without an increase in pHi, cells cannot complete the S/G2 transition but that dynamic pHi is required for correct S phase timing.

We did not measure a significant difference in G2 phase length with high pHi compared to CRL (Fig. 7G). Unfortunately, because low pHi cells could not complete the S/G2 transition, G2 phase times could not be measured for this treatment. M phase duration for low pHi cells could be measured only for cells in G2 or M during the start of the experiment. Based on these data, we hypothesized that if a high pHi threshold was already met during early G2, low pHi cells had the ability to complete division. This hypothesis aligns with the time-lapse data showing high pHi ∼5 h prior to division followed by a rapid acidification during mitosis (Fig. 4C, 5C). We saw no significant differences in M phase timing with pH manipulation. However, the longer acquisition window (20 minutes) of the time-lapse experiments in this work could also reduce accuracy of M phase length measurements.

The single-cell measurements presented here, both via endpoint assays and single-cell time-lapse measurements, show novel oscillating pHi dynamics throughout the cell cycle. Our data support prior work using ion transporter knockdown that showed high pHi regulates S phase length (Flinck et al., 2018a) and G2/M (Putney and Barber, 2003). However, our work also reveals novel decreases in pHi during G1/S, late S, and mitosis (Fig. 6A). Our combined use of single-cell pHi manipulation and cell cycle reporters show that pHi plays an important role in regulating the cell cycle, particularly for correct timing of G1 exit, S phase progression, and G2 entry (Fig. 8). Taken together, these results indicate that decreased pHi may be a cue for G1 exit but prevents cells from completing S/G2 and G2/M. We also found that dynamic increases/decreases are required for S phase, and increased pHi is necessary for G2 entry. In conclusion, our work suggests oscillating single-cell pHi not only reports on but regulates cell cycle progression in single cells.

**Figure 8:**
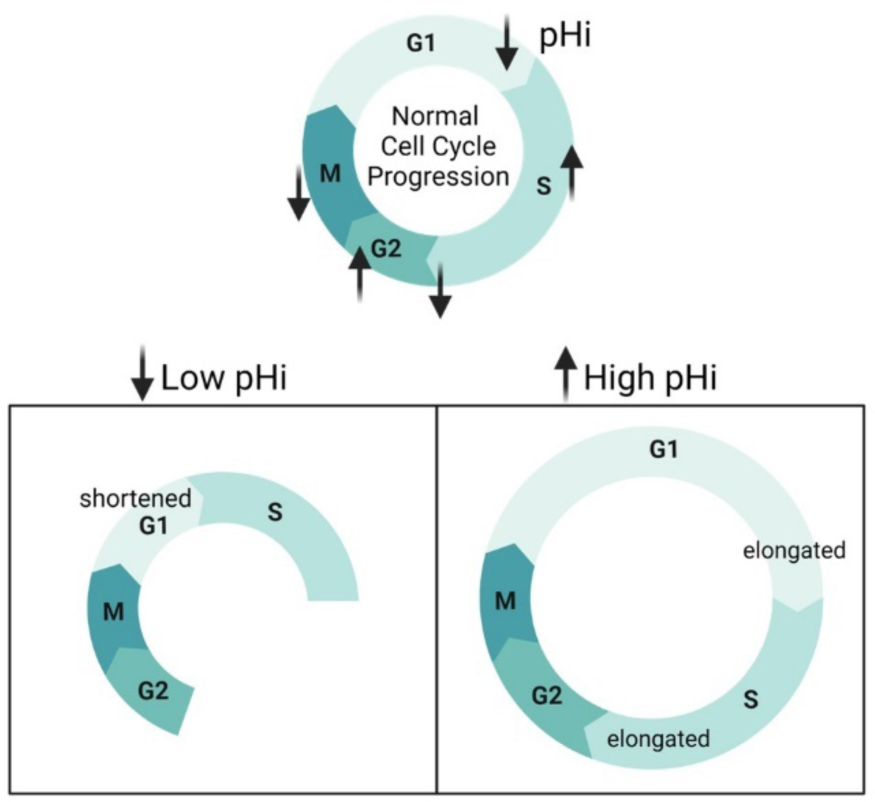
Single-cell pHi is dynamic during cell cycle progression and regulates G1 exit, S phase duration, and S/G2 transition. During cell cycle progression, pHi decreases at the G1/S boundary, increases in mid-S phase before dropping in late S, increases through G2 and decreases leading to division. When pHi is experimentally decreased, cells have a shortened G1 and less S/G2 transitions. When pHi is experimentally increased, G1 and S phases are elongated. This suggests that low pHi cues G1 exit and high pH is necessary for G2 entry.

## DISCUSSION

Intracellular pHi dynamics have been implicated in diverse cellular processes like differentiation (Ulmschneider et al., 2016), proliferation (Flinck et al., 2018b), migration (Martin et al., 2011), and apoptosis (Sergeeva et al., 2017). However, we have limited mechanistic understanding of how spatiotemporal and single-cell pHi dynamics regulate cell behaviors. This is partially due to reliance on population analyses, non-physiological environments, or genetic ion transporter ablation approaches to link pHi and phenotype (Czowski et al., 2020).

Here, we sought to characterize the relationship between pHi and cell cycle progression in single cells. Prior work at the population level suggested a role for pHi in regulating cell cycle progression (Flinck et al., 2018b). We show that single-cell pHi is dynamic and oscillates over an entire cell cycle, with pHi significantly decreasing at the G1/S boundary, increasing in mid-S, decreasing in late S phase, increasing through G2 and peaking at G2/M, before acidifying during mitosis and recovering in daughter cells. Here, we present three key results suggesting a regulatory link between pHi dynamics and cell cycle at both the population and single-cell levels.

First, we show that pHi significantly decreases at the G1/S boundary. These results were consistent regardless of which cell cycle synchronization method was used. The single-cell pHi manipulation data suggests that decreased pHi is a cue for G1 exit, as low pHi significantly shortened G1 and high pHi significantly elongated G1 compared to control cells. These results indicate a novel regulatory role for pHi acidification in regulating G1 exit. Future work will investigate what molecular drivers may be responding to the low pHi observed at the G1/S transition to time G1 exit or S phase entry in single cells.

Second, we show that pHi increases in mid-S phase and decreases before the S/G2 transition. Experimentally lowering pHi inhibited S/G2 transitions and raising pHi allowed for increased success of S/G2 transitions, but we also observed significantly elongated S phase with high pHi. These data suggest that increased pHi is necessary for successful entry into G2, but dynamic pHi (both increases and decreases) are important for proper timing of S phase. These data confirm prior results at the population level showing pHi increases in S phase (Flinck et al., 2018a) and successful transition through G2/M (Putney and Barber, 2003). However, our work also reveals novel temporally-regulated decreases in pHi that may be necessary for successfully timed G1/S and S/G2 transitions.

Third, we show that single-cell pHi peaks at G2/M and rapidly acidifies during M phase. The increase in pHi during G2/M confirms prior work showing increased pHi correlated with increased G2/M transition (Putney and Barber, 2003; Sellier et al., 2006). However, our data also suggests a novel role for intracellular acidification during late M phase and division. Supporting our results, recent work investigating intracellular lactate levels during cell cycle progression found that lactate regulates the anaphase promoting complex (APC) and is important for efficient mitotic exit (Liu et al., 2023). As lactate production releases protons as a byproduct, this recent work fits with the rapid acidification we measured in single cells during mitosis. Future work by our lab will apply optogenetic tools to spatiotemporally change pHi in single cells (Donahue et al., 2021) to further characterize roles for decreased pHi in regulating M phase timing or successful chromosomal segregation and division.

We also note that for most cells in the pHi manipulation experiments, we only monitored one division. Prior work in single-cell cell cycle progression has suggested that mother cell mitogen history affects daughter cell cycle (Min et al., 2020). Under E+S treatment, no daughter cells in the dataset successfully continued a second round of the cell cycle after the G1/S transition, though a handful of daughter cells divided again in the control population (∼25%). Future work will explore how dysregulated pHi dynamics in the mother cell alters or modulates daughter cell outcomes.

Prior work has suggested increased pHi in cancer promotes proliferation and tumorigenesis (Korenchan and Flavell, 2019; White et al., 2017a). Here, we show that while median pHi is increased in cancer cells compared to normal cells from the same tissue, single-cell pHi is heterogeneous and dynamic during cell cycle progression. However, the acidification of pHe also plays a role in maintaining these phenotypes (Boedtkjer and Pedersen, 2020) and altered pHe has been linked to cell cycle progression phenotypes in unicellular organisms (Aerts et al., 1985; Gillies and Deamer, 1979) as well as in mammalian cells (Deutsch et al., 1982). The 2D imaging in large relative volumes of bulk media performed here is unlikely to reflect proliferation in a tissue and extremely unlikely to mimic the competitive and confined environment that is produced during tumor growth and compression of surrounding tissue. Thus, in order to fully recapitulate roles for pHi in proliferation of a tissue or tumor, we must explore the role for spatiotemporal pHi dynamics in three dimensional environments. Future work will measure pHi gradients in normal and cancer cells in three dimensional environments with various extracellular matrix compositions and stiffnesses. This future work will have implications for how spatiotemporal pHi dynamics regulate biology and could lead to new therapeutic routes for limiting pHi-dependent behaviors in diseases with dysregulated pHi, such as cancer (Harguindey et al., 2017; White et al., 2017a) and neurodegeneration (Majdi et al., 2016).

Single-cell techniques can elucidate single-cell behaviors and reveal heterogeneity not found at the population level. Here, we addressed a critical need in the field to understand how pHi dynamics regulate single cells during cell cycle progression. These pHi dynamics could be essential for understanding complex cell biology that integrates single-cell and tissue-level behaviors. For example, prior work showed pHi gradients are generated in morphogenetic tissues (Weiß and Bohrmann, 2019). Our work now supports the hypothesis that bursts of synchronized cell proliferation may underlie these observations. More work is necessary to determine how temporal pHi gradients are generated during cell cycle phase transitions and whether a threshold of pHi changes is required. With our data establishing a framework of pHi regulation during an entire cell cycle, future work will determine which pH-sensitive proteins may be mediating and correctly timing pH-dependent cell cycle progression.

## MATERIALS AND METHODS

### Cell Culture and Conditions

Complete media for H1299 cells (ATCC CRL-5803): RPMI 1640 (Corning, 10-040-CV) supplemented with 10% Fetal Bovine Serum (FBS, Peak Serum, PS-FB2); A549 (ATCC CCL-185) and MDA-MB-231 (ATCC HTB-26): DMEM (Corning, MT10013CVV) supplemented with 10% FBS; and NL20 (ATCC CRL-2503): Ham’s F12 (Lonza, 12001-578) supplemented with 4% FBS, 1.5 g/L sodium bicarbonate (Sigma, S6014), 2.7 g/L glucose (VWR, BDH9230), 2.0 mM Glutamax (Gibco, 35050079), 0.1 mM nonessential amino acids (Lonza, BW13114E), 0.005 mg/mL insulin (Sigma, I1882), 10 ng/mL EGF (Peprotech, AF-100-15), 1 μg/mL transferrin (BioVision, 10835-642), 0.5 μg/mL hydrocortisone (Sigma, H0888). MCF10A (ATCC CRL-10317): DMEM/50% F12 w/ GlutaMax (Invitrogen, 10565-018) supplemented with 5% Horse Serum (Invitrogen, 16050-122), 0.02 μg/mL EGF (Peprotech, AF-100-15), 5 μg/mL Hydrocortisone (Sigma, H-0888), 0.01 mg/mL insulin (Sigma, I-1882), 0.1 μg/mL Cholera toxin (Sigma, C-8052), and 1% Penicillin-Streptomycin (Corning, 30-001-Cl). All cells were maintained at 5% CO_2_ and 37°C in a humidified incubator. All cell lines were authenticated and tested for mycoplasma in November 2022.

### Transfections and stable cell line selection

H1299 cells were transfected with the pCDNA3-mCherry-SEpHluorin (Koivusalo et al., 2010) (mCh-pHl) or pLenti-CMV-Blast-PIP-FUCCI using Lipofectamine2000 (Life Technologies, 11668019) per manufacturer’s instructions. After 24 h, cells were trypsinized and plated at a low dilution in a 10 cm dish with media containing 0.8 mg/mL Geneticin. Cloning cylinders were used to select colonies expressing mCh-pHl for expansion. A final clone was selected based on microscopy assay for mCh-pHl expression and comparison of cell morphology and pHi to parental H1299. For H1299-FUCCI, cells were trypsinized after 24 h transfection and plated at low dilutions (50 cells/mL) in a 96-well plate in media containing 0.8 mg/mL Blasticidin (Fisher, BP264725). Wells with equal expression were further expanded and screened on a microscopy assay for FUCCI expression and for similar cell morphology and pHi compared to parental H1299.

Lentiviral transfection was used to generate stable mCh-pHl expression in NL20 and A549 cells. Production of the virus was carried out in 293FT cells. Cells were grown to near confluency in a 10 cm dish and transfected with plx304-mCherry-SEpHluorin (Gift of Yi Liu and Diane Barber at UCSF) and two packaging plasmids: psPAX2 (Addgene #12260) and pmd2.G (Addgene #12259) provided by Siyuan Zhang (University of Notre Dame). Three µg each of the plx304-mCherry-pHluorin, psPAX2, and pmd2.G were transfected into a nearly confluent 10 cm dish of 293FT cells using Lipofectamine2000 for 18 h. Media was changed and incubated another three days. Viral supernatant was collected from the cells and centrifuged for 15 min at 3000 rpm. The supernatant was passed through a 0.2 µm polyethersulfone filter, flash-frozen in liquid nitrogen in 1 mL aliquots, and stored at −80°C.

NL20 and A549 cells were plated in a 6-well plate for viral transduction. After 24 h, viral supernatant was diluted 1:1.6, 1:3, and 1:10 into antibiotic-free media (depending on cell line) with 10 µg/mL Polybrene (Sigma, TR-1003-G), added to separate wells and incubated for 48-72 hr. Transduced cells were moved to a 10 cm dish and selected with 0.8 mg/mL Blasticidin. NL20 cells were plated at low density in 96-well plates (50 cells/mL). Colonies expressing mCh-pHl were expanded, and a final NL20-mCh-pHl clone was chosen with matched morphology and pHi of parentals. A549 cells were sorted using fluorescence-activated cell sorting (FACS), and a population sort according to mCherry expression was used for all imaging experiments after confirmation with microscopy.

### BCECF plate reader assays

Cells were plated at 4.0×10^5^ - 8.0×10^5^ cells/well in a 24-well plate and incubated overnight. Cells were treated with 2 μM 2’,7’-Bis-(2-Carboxyethyl)-5-(and-6)-Carboxyfluorescein, Acetoxymethyl Ester (BCECF-AM) (VWR, 89139-244) for 20 min at 37°C and 5% CO_2_. NL20 and H1299 cells were washed 3×5 minutes with a pre-warmed (37°C) HEPES-based wash buffer (30 mM HEPES pH 7.4, 145 mM NaCl, 5 mM KCl, 10 mM glucose, 1 mM MgSO_4_, 1 mM KHPO_4_, 2 mM CaCl_2_, pH 7.4) to match their low bicarbonate media (RPMI, Ham’s F12) and A549 cells were washed 3×5 minutes with a pre-warmed (37°C) Bicarbonate-based wash buffer (25 mM HCO_3_, 115 mM NaCl, 5 mM KCl, 10 mM glucose, 1 mM MgSO_4_, 1 mM KHPO_4_, 2 mM CaCl_2_, pH 7.4) to match its high bicarbonate media (DMEM). Two nigericin buffers (25 mM HEPES, 105 mM KCl, 1 mM MgCl_2_) were supplemented with 10 μM nigericin (Fisher, N1495), pH was adjusted to ∼6.7 and ∼7.7, and were pre-warmed to 37°C. Fluorescence was read (ex 440, em 535; ex 490, em 535) on a Cytation 5 (vendor: BioTek) plate reader incubated at 37°C with 5% CO_2_. Kinetic reads were taken at 15-sec intervals for 5 min, using a protocol established within Gen5 software. After the initial pHi read, the HEPES/bicarbonate wash was aspirated and replaced with one of the nigericin buffer standards, and cells were incubated at 37°C with 5% CO_2_ for 7 min. BCECF fluorescence was read by plate reader as above. This process was repeated with the second nigericin standard. As it takes significant time to equilibrate CO_2_ in the plate reader, we did not measure nigericin standardizations without CO_2_. The mean intensity ratio (490/440) was derived from each read. Measurements were calculated from a nigericin linear regression using exact nigericin buffer pH to the hundredths place (Grillo-Hill et al., 2014).

### Western Blot

Protein lysates were collected from 35-mm dishes or 6-well plates frozen at time points matched to imaging. Ice-cold lysis buffer [50 mM TRIS, 150 mM NaCl, 1 mM dithiothreitol (DTT), 1 mM Ethylenediaminetetraacetic acid (EDTA), 1% Triton X-100, Roche Protease Inhibitor Cocktail] was added to the samples and incubated for 15 min on ice. Cells were scraped and centrifuged for 10 min at 13,000 g at 4°C. The supernatant was retained, and protein concentration was determined by Pierce™ BCA (ThermoFisher, 23225) protein assay.

15 μg protein was loaded onto an SDS-Polyacrylamide gel electrophoresis (PAGE) that was run for 3 h at 120 V in 1X Tris-glycine (3.02 g/L Tris, 14.4 g/L glycine, 1.0 g/L SDS). Either a wet-transfer system or a Trans-Blot Turbo Transfer System (Bio-Rad) was used to transfer the proteins to a polyvinylidene fluoride (PDVF) membrane (pre-wet with methanol). For the wet transfer, 1X transfer buffer (141 g/L Glycine, 0.3 g/L Tris base) with 20% Methanol for 1.5 h at 100 V. For the Trans-Blot Turbo Transfer, Bio-Rad transfer buffer was used according to the manufacturer’s protocol (7 min). Membranes were blocked in 5% BSA in TBST (2.42 g/L Tris, 8 g/L NaCl, 0.1% Tween) for 2 h then divided for blotting. Primary antibodies: α-cyclin A2 (1:500; Abcam, ab38), α-cyclin B1 (1:1,000; Cell Signaling, 12231), α-cyclin E1 (1:1000; Cell Signaling, 4129), actin (1:1,000; Santa Cruz, 2Q1055). Membranes were incubated with primary antibody solution overnight at 4°C with shaking (4 h at RT with shaking for actin). Membranes were washed 3×10 min TBST at RT with shaking and incubated with secondary antibodies [1:10,000; goat α-mouse IgG HRP (Bio-Rad, 1721011) or goat α-rabbit IgG HRP (Bio-Rad, 1706515)] for 2 h at RT with shaking. Membranes were washed 3×10 min TBST at RT with shaking, developed using SuperSignal™ West Pico PLUS Chemiluminescent Substrate (ThermoFisher, 34578), and visualized using a ChemiDoc MP Imaging System (BioRad). ImageJ was used for protein quantification, normalized to loading control.

### Double-Thymidine block

Cells were plated at 10% confluency in 5 replicate 35-mm glass-bottomed dishes and 5 replicate 6-well plates (for protein lysate collection) and incubated overnight. Dishes were identically treated with 2 mM thymidine (Sigma, T9250) for 18 h, washed with PBS quickly (<30 sec), and incubated with fresh complete media for 9 h, then treated for another 18 h with 2 mM thymidine. Cells were released with a quick (<30 sec) PBS wash and replaced with fresh complete media. Imaging of the 0 h time point was initiated 20 min after release. Subsequent imaging was collected at 4, 8, 12, and 24 h after release in respective media. Matched dishes at each time point were washed twice with PBS and frozen at −80°C for protein lysate collection and immunoblot analysis of cyclins.

For time-lapse imaging, the double-thymidine block was used as explained above on a single 35-mm glass-bottomed dish supplemented with 1% Pen/Strep (Corning, 30-001-C1) to avoid bacterial contamination during time-lapse microscopy. Hoechst 33342 solution (ThermoFisher, 62249) was added to the cells (1:20,000) before release and incubated for 15 min. Dye and thymidine were removed, and cells were washed with PBS to release cells. Fresh media was added, and images were collected every 20 min for 24 hr. Acquisition parameters: 700 ms exposure time and 8% laser power for GFP; 700 ms exposure time and 10% laser power for TxRed; and 100 ms exposure time and 5% laser power DAPI. A single Z-plane was collected to avoid photobleaching. Nigericin standards were carried out as previously described (Grillo-Hill et al., 2014). For the asynchronous time-lapses, cells were plated the day prior to imaging and images were collected identically to thymidine-treated cells.

### Palbociclib Synchronization

Cells were plated at 10% confluency in 5 replicate 35-mm glass-bottomed dishes and 5 replicate 6-well plates (for protein lysate collection) and incubated overnight. Dishes were identically treated with 0.1 µM Palbociclib (PD-0332991) (Selleck, S1116) for 24 h. Cells were washed with PBS quickly (<30 sec) and released with complete fresh media. Imaging of the 0 h time point was initiated 20 min after release. Subsequent images were collected at 4, 8, 12, 24, and 36 h post-release in respective media (1 replicate 0-24 h, 2 replicates 0-36 h). Matched dishes at each time point were washed twice with PBS and frozen at −80°C for protein lysate collection and immunoblot analysis of cyclins.

### FUCCI cell cycle assays

For H1299-FUCCI time-lapses, cells were plated in a 4-well imaging dish (10,000 cells/well) and supplemented with 1% Pen/Strep (Corning, 30-001-C1) to avoid bacterial contamination during long-term acquisition. For HCO_3_^−^ supplementation, 35-mm glass-bottomed dishes were used. Hoechst dye was added to the cells (1:20,000) 2-4 h prior to imaging and incubated for 15 min, dye solution was removed, and fresh media was added to the cells. Experiments were started immediately after treatments were added to the cultured media and images were collected every 20 min for 36 hr. Optimal acquisition parameters were as follows: 200 ms exposure time and 8% laser power for GFP; 800 ms exposure time and 10% laser power for mCherry; and 200 ms exposure time and 5% laser power DAPI.

### Microscopy

Imaging protocol was derived from Grillo-Hill et al. Cells were plated on a 35-mm imaging dish with a 14-mm glass coverslip (Matsunami, D35-14-1.5-U) a day before imaging. Microscope objectives were preheated to 37°C, and the stage-top incubator was preheated to 37°C and kept at 5% CO_2_/95% air. Confocal images were collected on a Nikon Ti-2 spinning disk confocal with a 40x (CFI PLAN FLUOR NA1.3) oil immersion objective. The microscope is equipped with a stage-top incubator (Tokai Hit), a Yokogawa spinning disk confocal head (CSU-X1), four laser lines (405nm, 488nm, 561 nm, 647 nm), a Ti2-S-SE motorized stage, multi-point perfect focus system, and an Orca Flash 4.0 CMOS camera. Hoechst dye (405 laser ex, 455/50 em), pHluorin (488 laser ex, 525/36 em), TxRed (561 laser ex, 605/52 em), and mCherry (561 laser ex, 630/75 em), SNARF (561 laser ex, 705/72 em) were used. Acquisition times for each fluorescence acquisition ranged from 100-800 milliseconds.

### Immunofluorescence Assays

Cells were plated in a 4-well imaging dish (20,000 cells/well) overnight, then treated with pHi-manipulation media or etoposide (positive control to validate H2AX antibody, 10 μM) for 24 h. Cells were rinsed with PBS and fixed in 3.7% formaldehyde (Alfa Aestar, 33314) at room temperature for 10 minutes. Cells were washed 3×2 min with DPBS, then incubated lysing buffer (0.1% Triton-X in DPBS) for 10 minutes at room temperature. Cells were washed 3×2 min with DPBS, then incubated in blocking buffer (1.0% BSA in DPBS) for 1 hour at RT with rocking followed by 3×2 min wash in DPBS. Cells were then incubated overnight at 4°C with H2AX (1:100; Cell Signaling, 9718S) in antibody buffer (0.1% Triton-X, 1.0% BSA in DPBS). Cells were washed 3×2 min in DPBS, then incubated with secondary antibody (Alexa Fluor 488 goat anti-rabbit, 1:1000, Invitrogen, A11008) for 1 h at room temperature. After 3×2 min washes in PBS, Hoechst dye (1:20,000) in antibody buffer was added for 15 min then removed. Cells were imaged in DPBS on a spinning disk confocal previously mentioned: Hoechst dye (405 laser ex, 455/50 em), AlexaFlour488 (488 laser ex, 525/36 em).

### SNARF microscopy assays

We note that we were not able to isolate an mCherry-pHluorin stable cell line for MCF10A, so comparisons between MCF10A and MDA-MB-231 were performed using the pH-sensitive dye SNARF [4-(and-6)-CarboxySNARF-1 acetoxymethyl ester, acetate] (Invitrogen, C1272). Cells were plated at 4.0×10^5^ cells/well in an imaging dish (Matsunami). Conditioned media was removed and cells were treated with 20 μM SNARF in serum-free media for 15 minutes and then media was replaced with conditioned media. Images were collected similarly to mCh-pHl experiments using nigericin standardization.

### Intracellular pH imaging and data collection

For all pHi imaging (SNARF and mCherry-pHluorin), initial fields of view (FOV) were collected on the cells in their respective media. For all imaging, nigericin buffers were prepared identically to BCECF assays, and all buffer exchanges were carried out on the stage incubator to preserve XY positioning. Multiple Z-planes were collected with the center focal plane maintained using the Nikon Ti2 Perfect Focus System (PFS).

For time-lapse pHi and FUCCI imaging, a single Z-plane was collected to avoid excess light and additional water was added to the stage top incubator at 18 h. For pHi manipulation validation, cells were plated at 20% confluency on a 35-mm imaging dish with a 14-mm glass coverslip and incubated overnight. For Fig. 6, cells were treated with a combination of 15 μM EIPA/30 μM S0859 (E+S), 1 µM Concanamycin A (CMA), 100 mM NaHCO_3_^−^ (HCO_3_^−^), or 20 mM ammonium chloride (NH_4_Cl) diluted in fresh media and incubated for 24 h. For Fig. S8, cells were treated with E+S or NH_4_Cl for 4, 8, or 12 h. Both imaging collection and pHi calculations were completed identically to other single-cell pHi measurement experiments. E+S, HCO_3_^−^, and NH_4_Cl treated cells with respective controls were corrected for photobleaching by collecting images of cells in nigericin buffers (pH 7.4) with treatment supplemented but without nigericin present.

### Image Quantification

Images were background-subtracted using an ROI drawn on a glass coverslip (determined by DIC). For pHi quantification, individual Regions of Interest (ROI) were drawn for each cell in each condition (initial, high pH nigericin, and low pH nigericin). For SNARF assays, mean TxRed and SNARF pixel intensities were quantified for each cell and SNARF/TxRed ratios were calculated in excel. For mCherry-pHluorin assays, mCherry aggregates are removed using thresholding holes and then each pHluorin and mCherry pixel intensities were quantified for each cell. The pHluorin/mCherry ratios were calculated in excel. In both cases, a cutoff of 100 a.u. was used for both pHluorin and mCherry intensity values after exporting. For each cell, nigericin values were used to generate a standard curve, and pHi was back-calculated from the single-cell standard curve.

For FUCCI analysis, cells were tracked using NIS Analysis software and nuclear regions of interest (ROI) based on DNA stain. In case of improper tracking, manual tracking was used to redraw ROIs. Manual tracking was also used during mitosis when the signals diffused throughout the cell. mVenus and mCherry intensities were exported from matched single-cell nuclear ROI at each time point over 36 h. Cell cycle phases were determined by mVenus or mCherry fluorescence intensity, adapted from Grant et al. For each individual cell trace, including subsequent daughter cells, an excel macro was used to determine timepoints for mVenus and mCherry cutoffs. G1/S was defined as a decrease in mVenus signal below 5% of maximum mVenus intensity. As validation of G1/S, S phase entry was defined as the first time point after mCherry minimum that showed a 3% (determined from mCherry maximum) increase in mCherry intensity compared to the previous point. S/G2 was defined as point at which mVenus intensity rose above 2% of mVenus maximum compared to the previous point. G2/M and M/G1 were defined by nuclear envelope breakdown and division into two daughter cells, respectively.

For H2AX staining, after background subtraction, the nuclear regions of interest (ROI) were drawn based on DNA stain and GFP intensities were exported for each treatment condition.

### Statistics

GraphPad Prism was used to prepare graphs and perform statistical analyses. Normality tests were performed on all data sets as well as outlier test using the ROUT method (Q=1%). For normally distributed data, an unpaired t-test (Fig. S1A-C; Fig. 5G) or paired t-test (Fig. 2F-H; Fig. 3F-H) was used. A Mann Whitney test was used for non-normal, unpaired data (Fig. 1F-G; Fig. S3C; Fig. S4C; Fig. S5F; Fig. 6B, 7E-H; Fig. S8). For time-lapse data (Fig. 4C, 4F; Fig. 5C, 5F), a one-sample Wilcoxon test was used, compared to a theoretical mean of 0. For non-normal, unpaired data with more than two sets, a Kruskal-Wallis test with Dunn’s multiple comparisons test was used (Fig. 2D, 2I; Fig. S3A; Fig. 3D, 3I; Fig. S4A; Fig. S5C, S5E). Values were binned at 0.02 in all frequency distributions. All significance was indicated in figures by the following: *p<0.05; **p<0.01; ***p<0.001.

## ACKNOWLEDGEMENTS

We would like to thank Dr. Siyuan Zhang (University of Notre Dame) and Dr. Diane Barber (University of California San Francisco) for plasmids. We would also like to thank members of the White lab for their helpful feedback on the manuscript. Figure schematics (Fig. 1A, 2A, 3A, 5D, 6A, 8) created with BioRender.com.

## AUTHOR CONTRIBUTIONS

JSS & KAW Conception and design. JSS & KAW acquisition of data. JSS & KAW acquisition, analysis, and interpretation of data; drafting and revising manuscript text.

