## Supplemental Figures for "Single-cell intracellular pH dynamics regulate the cell cycle by timing G1 exit and the G2 transition"

**SUPPLEMENTAL INFORMATION (SI)**

**SUPPLEMENTAL VIDEO LEGENDS:**

**Movie 1: Single, dividing cell from an asynchronous population shows dynamic pHi changes during cell cycle progression.** Video of a dividing cell in an asynchronous population over 24 h, compiled from a single Z-plane captured every 20 min. Arrows show cell of interest and reappear to show timing of prophase/cytokinesis and two daughter cells, scale bar: 20 µm. Ratiometric display of pHluorin/mCherry fluorescence ratios. Video is the same cell as Fig. 4A and ratiometric scale is identical.

**Movie 2: Single, non-dividing cell shows attenuated pHi dynamics over a 24-hour period.** Video of a non-dividing cell from an asynchronous population over 24 h, compiled from a single Z-plane captured every 20 min. Arrows show cell of interest at the beginning and end of video, scale bar: 20 µm. Ratiometric display of pHluorin/mCherry fluorescence ratios. Video is the same cell as Fig. 4D and ratiometric scale is identical.

**Movie 3: Single, dividing cell released from S phase synchronization shows dynamic pHi changes during cell cycle progression.** Video of a dividing cell over 24 h after release from a double thymidine block, compiled from a single Z-plane captured every 20 min. Arrows show cell of interest and reappear to show timing of prophase/cytokinesis and two daughter cells, scale bar: 20 µm. Ratiometric display of pHluorin/mCherry fluorescence ratios. Video is the same cell as Fig. 5A and ratiometric scale is identical.

**Movie 4: Single, non-dividing cell shows attenuated pHi dynamics over a 24-hour period.** Video of a non-dividing cell over 24 h after release from a double thymidine block, compiled from a single Z-plane captured every 20 min. Arrows appear to show cell of interest at the beginning and end of video, scale bar: 20 µm. Ratiometric display of pHluorin/mCherry fluorescence ratios. Video is the same cell as Fig. 5D and ratiometric scale is identical.

**Movie 5: PIP-FUCCI accurately reports cell cycle phases in H1299 cells.** Video of a dividing H1299-FUCCI cell over 24 h (20 min intervals), compiled from a single Z-plane captured every 20 min. PIP-FUCCI uses regulatory domains PCNA interacting protein degron from human $Cdt1_{1-17}$ (PIP) (mVenus, green) and $Geminin_{1-110}$ (mCherry, magenta) to differentially express fluorescent proteins during cell cycle phases. Labels appear as the cell transitions to each phase (G1, S, G2, M, G1), scale bar: 20 µm. PIP-mVenus accumulates in the nucleus during G1 and is rapidly lost during onset of S phase, where mCherry-Geminin begins to rise and is expressed throughout S, G2, and M phases. During the S-G2 transition, mVenus accumulates again, and both mVenus and mCherry are co-expressed until division. M phase is marked by nuclear envelope breakdown and diffusion of mVenus and mCherry signals throughout the cell. Following cytokinesis, only mVenus is expressed in the two daughter cell nuclei, marking G1. Video is the same cell as Fig. 6C.

### SUPPLEMENTAL FIGURES

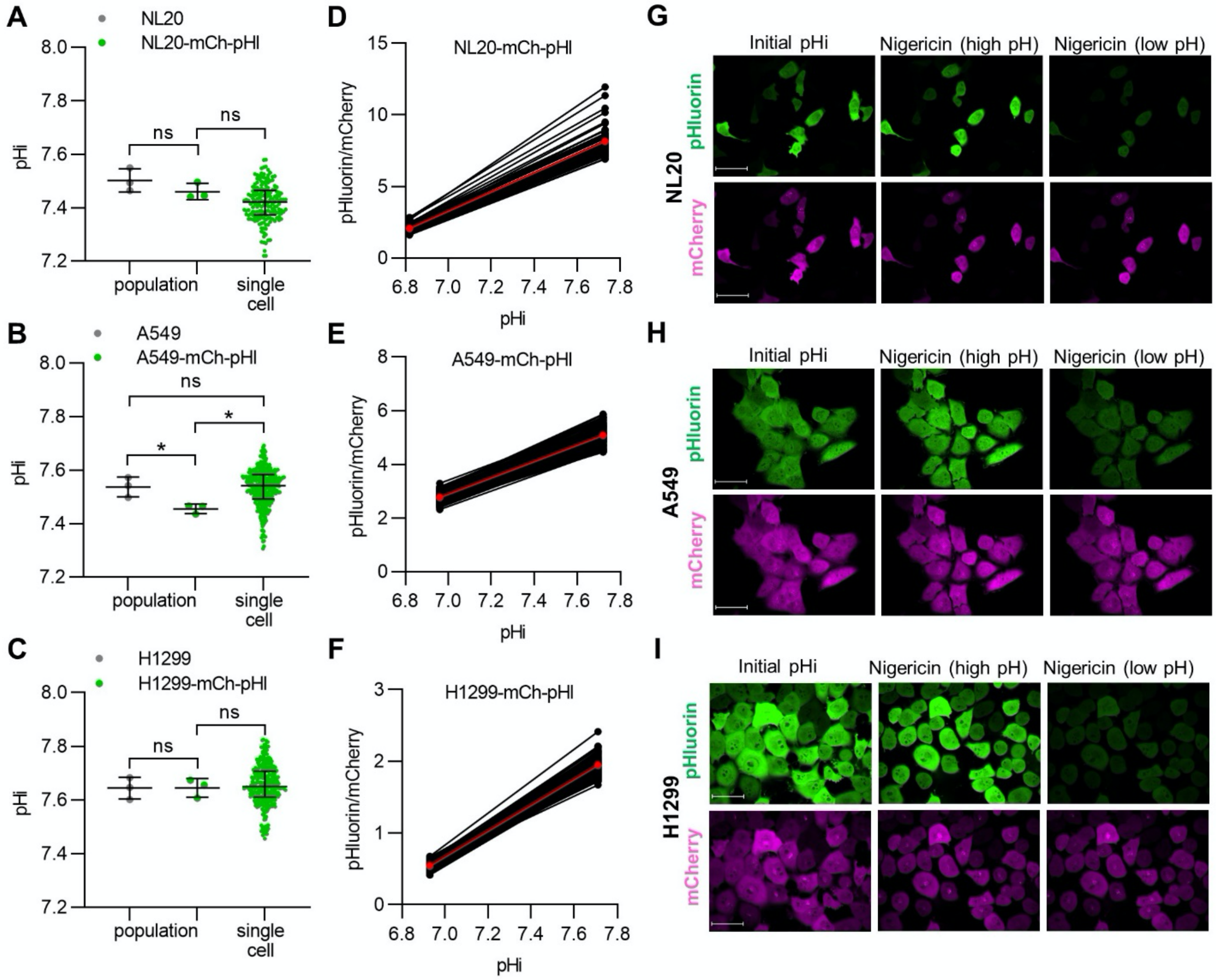


**Figure S1: Stable biosensor expression does not alter intracellular pH (pHi) compared to parental cell lines and example ratiometric quantification of mCherry-pHluorin.** Population-level plate reader assays (see methods) were performed on A) NL20 parental (n=3, 7.50±0.04) and NL20-mCh-pHI cells (n=3, 7.46±0.03); B) A549 parental (n=3, 7.54±0.04) and A549-mCh-pHI cells (n=3, 7.46±0.02); and C) H1299 parental (n=3, 7.64±0.04) and H1299-mCh-pHI cells (n=3, 7.65±0.03) with single-cell data of mCh-pHI lines reproduced from Fig. 1 for comparison. For A-C BCECF population data, unpaired t-tests (two-tailed) were performed. (*$p<0.05$). Single cell data was not significantly different from population pHi for NL20-mCh-pHI ($p=0.292$) and H1299-mCh-pHI ($p=0.825$). For A549 cells, pHi measurements from A549 mChpHI population assays was significantly lower ($p=0.0266$) than parental A549, but not and single-cell A549-mCh-pHI data showed no significant difference in pHi compared to parental A549 ($p=0.938$). Nigericin standard linear regressions across a single replicate for each single cell (black) and mean (red) for D) NL20-mCh-pHI, E) A549-mCh-pHI, and F) H1299-mCh-pHI. Single channel images of pHluorin (green) and mCherry (magenta) fluorescence at initial pHi imaging, high nigericin, and low nigericin standardizations for G) NL20-mCh-pHI, H) A549-mCh-pHI, and I) H1299-mCh-pHI. LUTs are identical across each cell line. Note that in all cases, single-cell nigericin standard curves were used to back-calculate single-cell pHi. Means are shown here simply for ease of comparison across the cell lines.

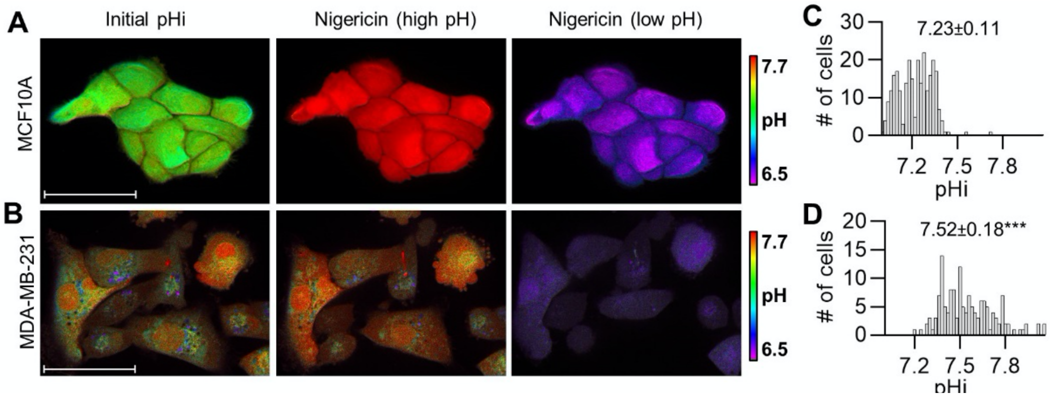


**Figure S2: Triple negative breast cancer cells have significantly increased pHi compared to normal breast epithelial.**

Representative images of pHi measurements and standardization in A) normal breast epithelial (MCF10A) and B) triple-negative breast cancer (MDA-MB-231) cells loaded with the pH sensitive dye SNARF (see methods). Ratiometric display of 640/580 emission fluorescence ratios, scale bars: 50 µm. B) Histograms of single-cell pHi in C) MCF10A cells (n=264, 3 biological replicates) and D) MDA-MB-231 cells (n=147, 3 biological replicates). Histogram is binned at 0.02 pH units. Above histogram, median±S.D is shown. Significance was determined by a Mann Whitney test. (***$p<0.001$, compared to MCF10A).

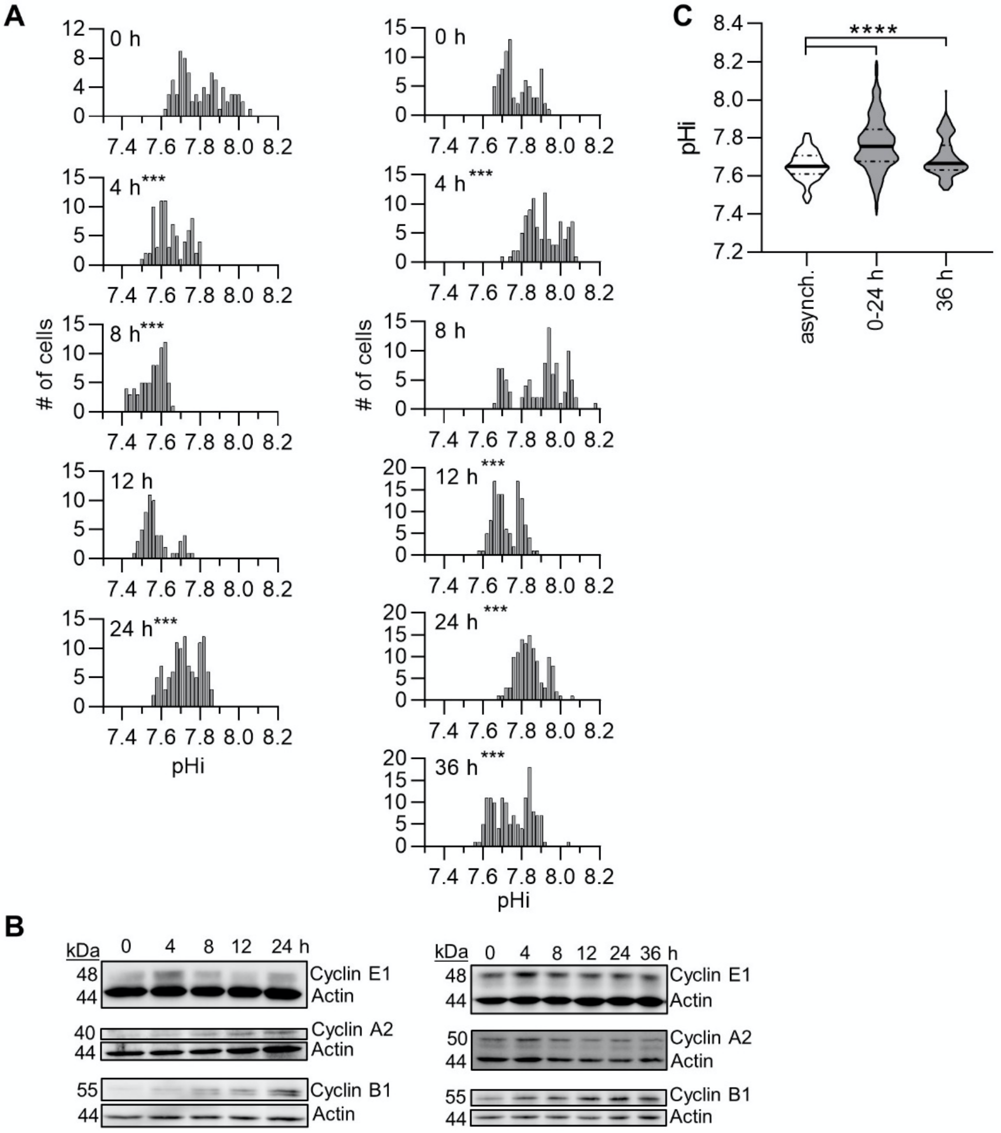


**Figure S3: Intracellular pH is dynamic after release from G1 in H1299-mCh-pHI cells and correlates with cyclin levels (Additional Replicates; data summarized in Figure 2I).** A) Histograms of single-cell pHi data after Palbociclib release, collected from 2 biological replicates. Histograms binned at 0.02 pH units. B) Representative immunoblots for cyclin E1, A2, and B1 with respective actin loading controls. C) Median pHi of Palbociclib-treated cells, both synchronized (0-24 h) and unsynchronized (36 h), compared to untreated, asynchronous (asynch.) cells collected in Fig. 1G. For A, significance was determined using a Kruskal Wallis test with Dunn's multiple comparisons correction. Each time point was compared to its preceding time point. For C, significance was determined by a Mann Whitney test. (***p<0.001).

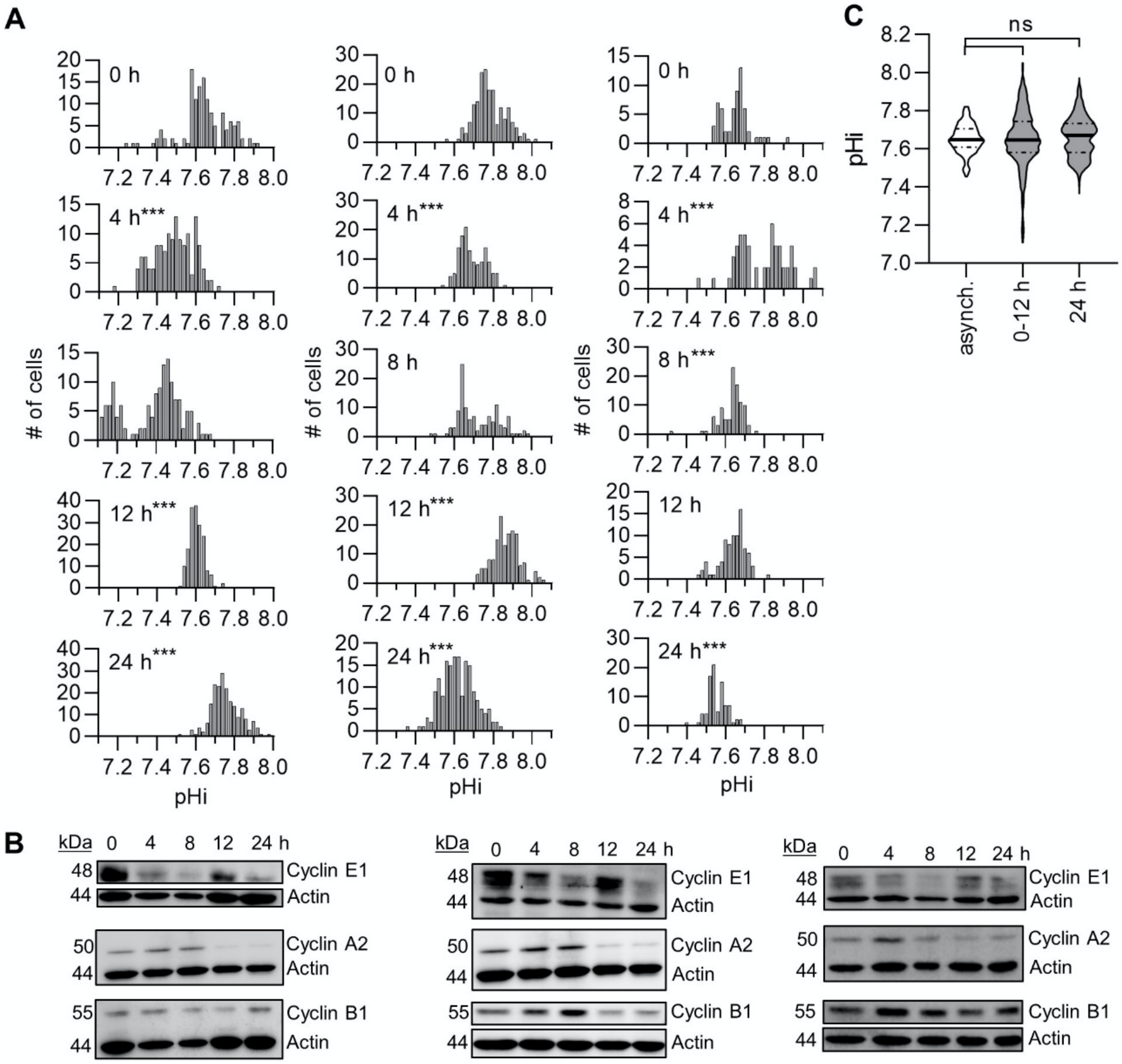


**Figure S4: Intracellular pH is dynamic after release from early S in H1299-mCh-pHI cells and correlates with cyclin levels. (Additional Replicates; data summarized in Figure 3I).** A) Histograms of single-cell pHi measurements after release from a double-thymidine block, collected from 3 biological replicates. Histograms binned at 0.02 pH units. B) Representative immunoblots for cyclin E1, A2, and B1 with respective actin loading controls. C) Median pHi of thymidine-treated cells, both synchronized (0-12 h) and unsynchronized (24 h), compared to untreated, asynchronous (asynch.) data collected in Fig. 1G. For A, significance was determined using a Kruskal Wallis test with Dunn's multiple comparisons correction. Each time point was compared to its preceding time point. (***$p<0.001$). For C, significance was determined by a Mann Whitney test.

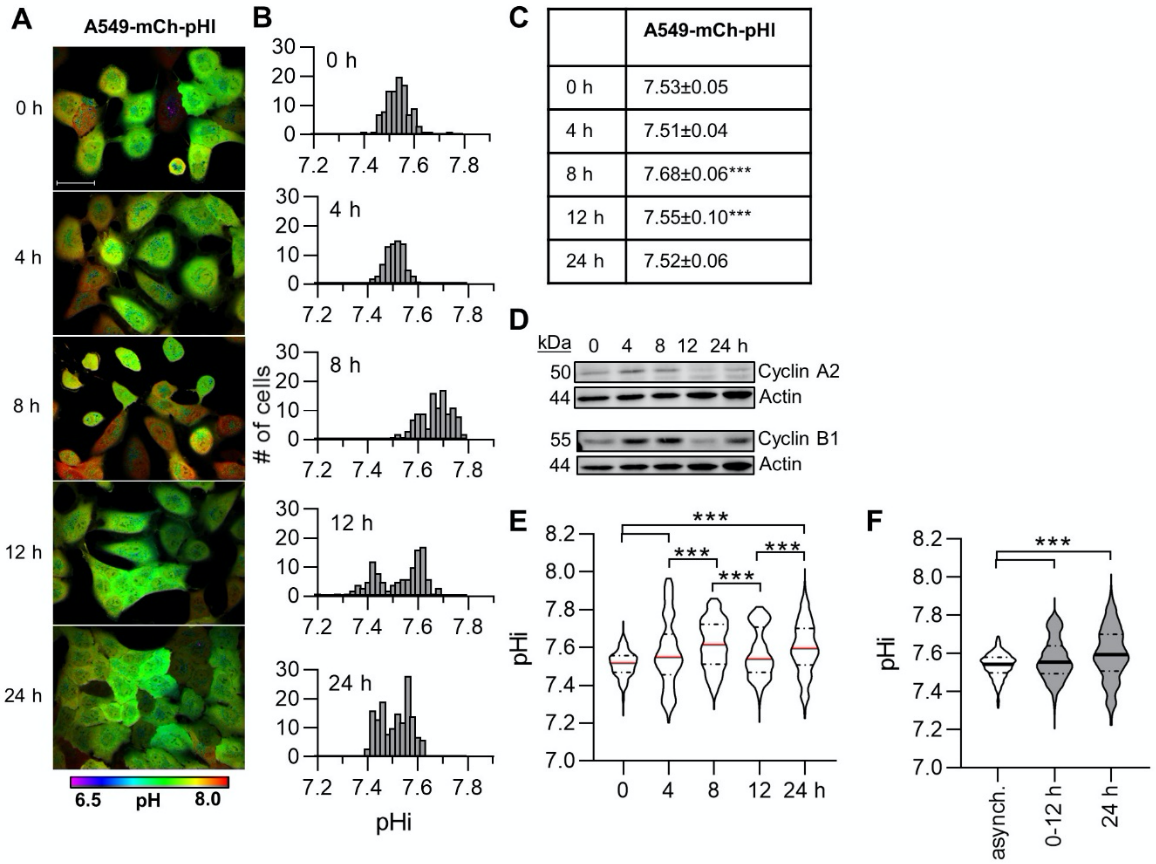


**Figure S5: Intracellular pH is dynamic after release from early S in A549-mCh-pHl cells and correlates with cyclin levels**. A) Representative images of A549-mCh-pHl cells at indicated time points after release from a double-thymidine block. Ratiometric display of pHluorin/mCherry fluorescence ratios, scale bars: 50 µm. B) Histograms of single-cell pHi data collected as in A, from one biological replicate. Histograms binned at 0.02 pH units. C) Table of pHi values from data in B, (median±S.D.). D) Representative immunoblots for cyclin E1, A2, and B1 with respective actin loading controls. E) Violin plots of raw pHi across 4 biological replicates. F) Median pHi of thymidine-treated cells, both synchronized (0-12 h) and unsynchronized (24 h), compared to untreated, asynchronous (asynch.) data collected in Fig. 1F. For C and E, significance was determined using a Kruskal Wallis test with Dunn's multiple comparisons correction. Each time point was compared to the preceding time point and 0 h compared to 24 h. (***$p<0.001$). For F, significance was determined by a Mann Whitney test. (*$p<0.05$; **$p<0.01$; ***$p<0.001$).

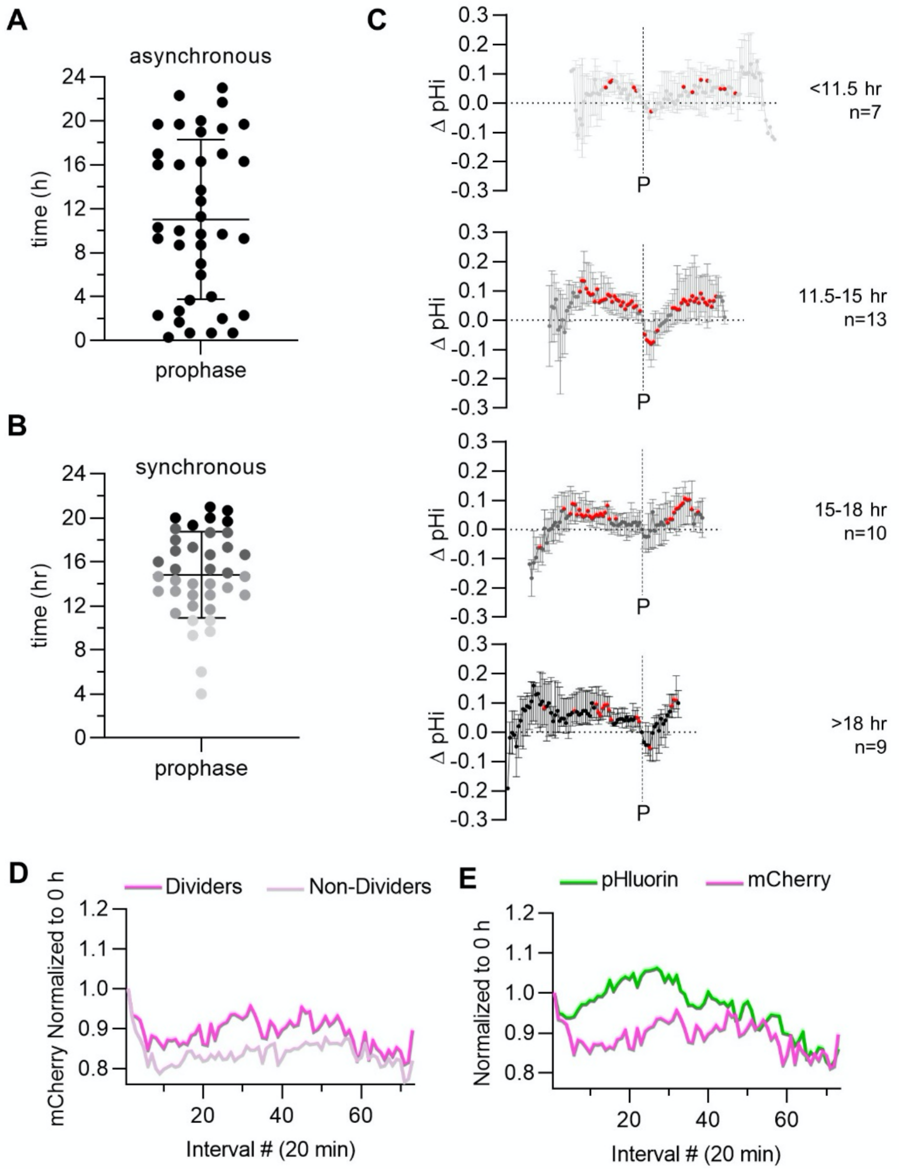


**Figure S6: Cells released from early S synchronization show bursts of division and pHi dynamics are preserved independent of mitosis timing. Normalized biosensor fluorescence intensities indicate pHluorin changes not an artifact of biosensor expression level changes.** A) Timing of prophase in asynchronous cell populations (n=39, 4 biological replicates). B) Timing of prophase after release from a double-thymidine block, where different shades of gray represent prophase bins (<11.5 h, 11.5-15 h, 15-18 h, >18 h) (n=39, 3 biological replicates). C) Change in pHi from prophase (P) for each individual cell released from S phase synchronization, binned according to timing of prophase in B (median±IQ range). In C, significance is determined by one-sample Wilcoxon test compared to 0 (red points, *p<0.05). D) Median mCherry intensity values, normalized to the intensity at 0 h for each cell, for dividing (dark pink) and non-dividing (light pink) cells in thymidine-synchronized cells. The x-axis shows time interval number (20 min intervals). E) Median mCherry and pHluorin intensity values for thymidine-synchronized cells. The x-axes show time interval number (20 min intervals).

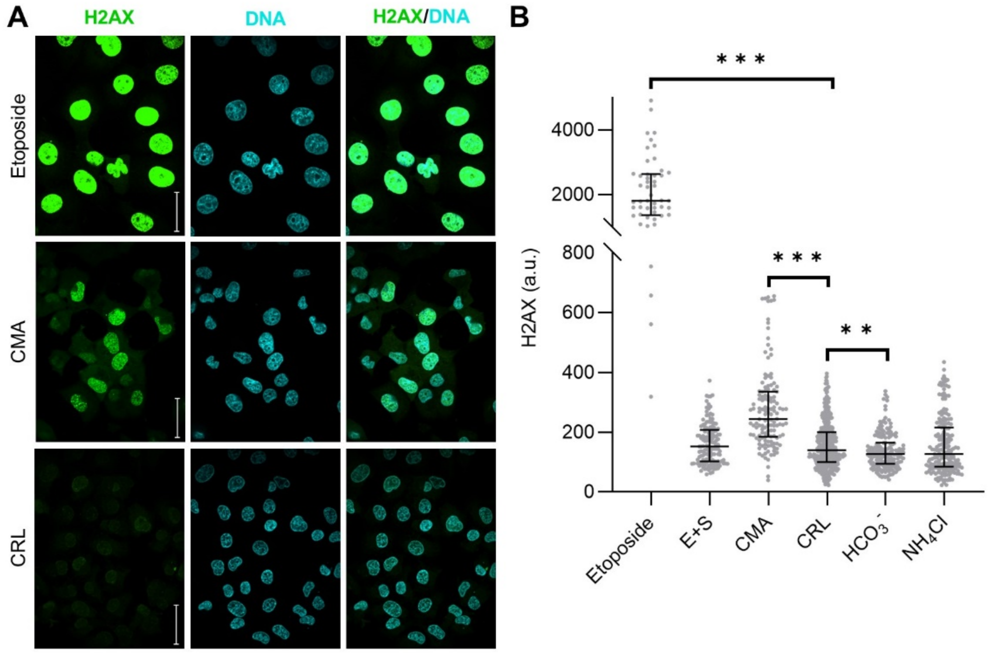


**Figure S7: Concanamycin A (CMA) treatment significantly increased gamma-H2A histone family member X (H2AX) foci, a marker of replicative stress.** A) Representative images of H2AX (green) staining in cells treated with double-strand break inducing Etoposide (10 µM), CMA (1 µM), or untreated (CRL) cells. Cells were co-stained with Hoescht dye (DNA, cyan) and overlay of H2AX/DAPI is shown, scale bars: 50 µm. B) Scatter plots of H2AX intensity in single cells treated with Etoposide or other pHi manipulation techniques shown in Fig 6B (15 µM EIPA + 30 µM SO859; 1 µM CMA; 100 mM $NaHCO_3$; and 20 mM $NH_4Cl$). Significance determined by an unpaired t-test compared to CRL (***p<0.001, **p<0.01).

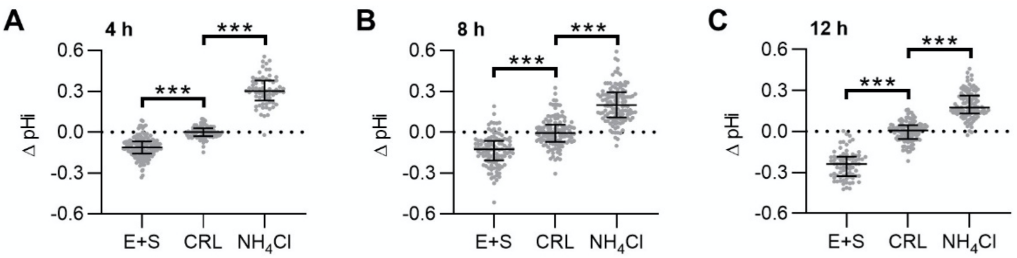


**Figure S8: Intracellular pH is significantly altered within 4 h of pH manipulation.** Single-cell pHi of H1299-mCh-pHl cells after A) 4 h, B) 8 h, and C) 12 h treatment with 15 μM EIPA and SO859 30 μM (E+S), untreated (CRL), and 20 mM ammonium chloride ($NH_4Cl$) A) 4 h (E+S, n=176; CRL, n=88; $NH_4Cl$, n=65) B) 8 h (E+S, n=110; CRL, n=120; $NH_4Cl$, n=124), C) 12 h (E+S, n=76; CRL, n=105; $NH_4Cl$, n=99). (2 biological replicates; 12 h CRL and $NH_4Cl$, 3 biological replicates). Scatter plot (median±IQ range). Mann-Whitney test to determine statistical significance (***p<0.001).
